# Predicting Mouse Lifespan-Extending Chemical Compounds with Machine Learning

**DOI:** 10.1101/2024.10.29.620854

**Authors:** Aleksey V. Belikov, Caio Ribeiro, Christopher K. Farmer, Michael Petrascheck, João Pedro de Magalhães, Alex A. Freitas

## Abstract

Pharmacological interventions targeting the biological processes of ageing hold significant potential to extend healthspan and promote longevity. This, to our knowledge, is the first study that uses Machine Learning models trained specifically on mouse lifespan data (from DrugAge) to predict lifespan-extending compounds. The use of mammalian data significantly elevates translational relevance compared to previously available models trained predominately on *C. elegans* data. Our most successful Random Forest classifiers were trained on direct drug-target annotations, including Gene Ontology, UniProt Keywords, pathways (KEGG, Reactome, Wiki) and protein domains (InterPro), whereas models trained on gene expression (LINCS) and chemical substructures (PubChem) underperformed. Models trained on male datasets performed better than those trained on mixed-sex and female datasets, with the latter suffering from severe class imbalance due to much fewer positive-class instances. Notably, features related to G-protein coupled receptors, especially receptors for neurotransmitters, metabolic hormones and sex hormones, were identified as strong predictors of lifespan extension. We used ensemble classifiers comprised of top models to screen compounds from DrugBank, highlighting novel candidates for longevity studies. Major clusters of compounds with the highest predicted longevity-promoting effects target IGF1 and insulin receptors, beta adrenergic receptors, carbonic anhydrases, dopamine and serotonin receptors, voltage-gated potassium and calcium channels, sodium-dependent dopamine, serotonin and noradrenalin transporters, muscarinic acetylcholine receptors and adenosine receptors. We tested 22 predicted compounds in C. elegans and found that 6 of them significantly extended median lifespan: dihydroergotamine, mianserin, bromocriptine, voxtalisib, bms-754807 and solifenacine. We have also created a public web server with our top performing classifier ensembles: https://www.cs.kent.ac.uk/projects/lodprime/ Our study not only provides an important contribution to the longevity pharmacology field but also informs research on the fundamental mechanisms of ageing.

## 1 Introduction

The use of pharmacological interventions to promote healthspan and longevity, and to target the biological process of ageing, is currently a very active research area ^1–4^. Hundreds of drug treatments have been successfully applied to significantly increase the lifespan of model organisms such as *Caenorhabditis elegans* and *Mus musculus*^2,3^.

The large amounts of data related to developing longevity-promoting treatments can be analysed through the use of machine learning (ML) tools such as classification models, which are first trained on a set of instances (examples) and then are able to make predictions about previously unseen data^5^. In addition to making predictions, classification models can be analysed to give researchers insights into the prediction problem, such as predictive patterns shared by groups of instances (chemical compounds) or ranking the predictive features by their relative importance for classifying instances. The results from machine learning experiments can inform *in vivo* experiments and help guide the development of new drugs or treatments.

Previous studies that focus on other model organisms, such as *Caenorhabditis elegans*, have applied machine learning to longevity data ^6–17^. However, to the best of our knowledge, this is the first work that applies classification models from machine learning to predict whether a chemical compound promotes longevity specifically in mice. Using *Mus musculus* as the target organism is more challenging than using simpler organisms, mainly due to high cost and hence the limited number of lifespan experiments that can be used for training the models. However, compared to non-mammal model organisms, the insights into the biological mechanisms of ageing that apply to mice are significantly more likely to apply to other mammals such as humans ^18–20^. In most countries, mice remain a mandatory preclinical step in drug discovery pipelines.

In order to create the datasets to train our classification models, we obtained the mouse data from the DrugAge database ^21–23^, which compiles results of peer-reviewed research on lifespan extension in model organisms, including the results from the Interventions Testing Program (ITP) ^24^ (Figure 1). Each compound-sex combination is assigned a class label (positive/negative) that indicates whether it is associated with lifespan extension in mouse experiments.

**Figure 1.**
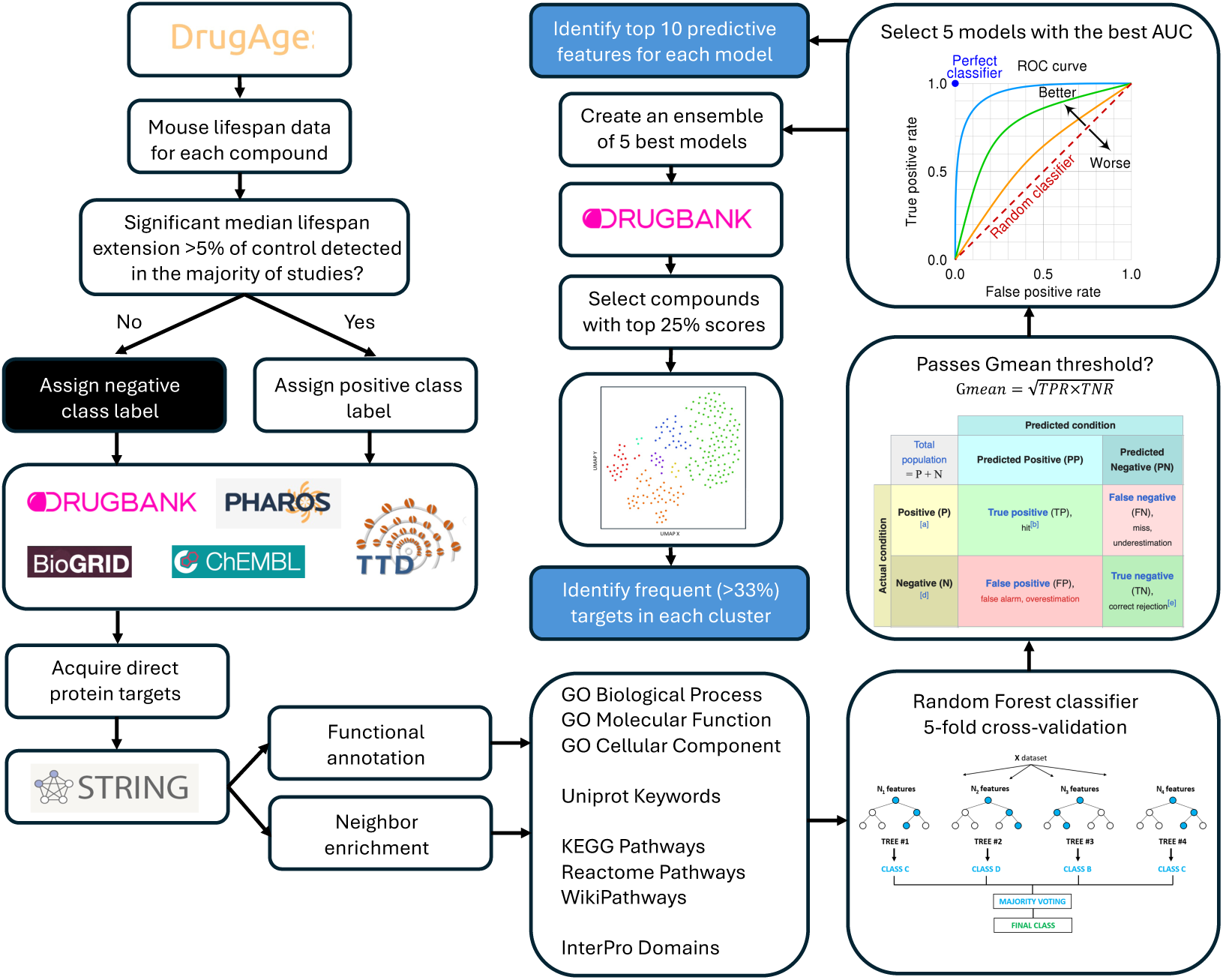
The study pipeline. Mouse lifespan data for each compound was acquired from DrugAge. If significant median lifespan extension above 5% of control was detected in the majority of studies, the compound was assigned a positive class label, otherwise it was assigned a negative class label. Direct protein targets of these compounds were acquired from DRUGBANK, PHAROS, BioGRID, ChEMBL and TTD. Functional annotation and neighbour enrichment of these targets were performed in STRING. The resulting GO, Uniprot, pathway and domain labels were used as features for training separate random forest classifiers with 5-fold cross-validation. Out of classifiers that passed G-mean threshold, we selected 5 models with the best AUC and identified top 10 predictive features for each model. We then combined 5 best models into an ensemble and applied it to the whole DRUGBANK dataset to assign a lifespan extension probability score to each compound. We then selected compounds with top 25% scores, clustered them using UMAP and identified frequent targets in each cluster, i.e. targets of more than 33% of drugs in that cluster.

Naturally, the efficacy of ML predictive models depends largely on the data used to train them. As longevity experiments vary in treatment regimens, mouse strains and lifespan metrics, it is more difficult for the predictive models to make correct generalizations, which are required for making accurate predictions on unseen data. One of the objectives of our study was to explore several different feature types and form a varied baseline of datasets. We prepared multiple datasets for training machine learning models, using three data sources, namely STRING^25^ (protein-protein interaction networks and protein annotations), LINCS^26^ (gene expression data) and PubChem^27^ (chemical substructure data). The data sources contain several principally different types of features that represent biological and chemical information about each compound such as the Gene Ontology, protein domains and pathway annotations of their targets. By preparing datasets that use a wide array of descriptors, we were able to explore the efficacy of orthogonal approaches for the task of predicting longevity-related compounds for mice.

We evaluated the predictive accuracy of classification models trained with each dataset and selected 10 models (5 from mixed-sex datasets and 5 from male-only datasets) to analyse further. We then identified features deemed most relevant for predicting whether a compound belongs to the positive class (promotes mice longevity). Models learned from female-only datasets were not analysed in detail because they led to smaller predictive accuracies than male-only or mixed-sex datasets.

In addition to the feature importance analysis, we looked into the negative-class compounds (based on DrugAge data) that were consistently classified as belonging to the positive-class (false-positive classifications) by our selected models. We believe these compounds, e.g. LY444711, putrescine, chlorpheniramine, and dehydroepiandrosterone sulphate, can be promising options for future longevity studies in mice, as although previous results did not find significant life extension in treatments with them, they share important characteristics with current successful life extension compounds.

We then combined the selected models into male-only and mixed-sex ensembles of classifiers and used these ensembles to classify thousands of previously unseen and unlabelled instances from an external database – DrugBank ^28^. We then clustered the top 25% predicted compounds with the highest predicted probability of positive class according to their feature profile similarity and identified the most common protein targets in each cluster. We also highlighted compounds acting on the target in the same way as positive compounds in the training dataset as promising options for future longevity studies.

Finally, we selected 22 most promising compounds from those predicted by our ensembles and subjected them to lifespan testing in *C. elegans*. We identified 6 compounds as significantly extending median lifespan: dihydroergotamine, mianserin, bromocriptine, voxtalisib, bms-754807 and solifenacine. It is important to note that many compounds that extend lifespan in mice (e.g. in ITP^29^) do not extend lifespan in *C. elegans* (e.g. in *Caenorhabditis* Intervention Testing Program (CITP)^30^) and vice versa^31^. Thus, the validity of our predictions is not limited by the *C. elegans* results, and multiple other compounds from our list might extend mouse (and human) lifespan. However, since our models were trained on mouse data, the compounds which also extended lifespan in worms are likely to act on evolutionary conserved longevity pathways, including the IGF-1R/IR-PI3K-mTor pathway^32^.

## Methods

### 1.1 Instance definition and labelling for machine learning

Each instance in our datasets represents a combination of a compound and the sex of the mice used in the experiments, based on data obtained from DrugAge build 5 ^21–23^, a database that collates results from peer-reviewed studies on the effects of chemical compounds (drugs) on the lifespan of model organisms. Its *Mus musculus* section contains data on more than a hundred different compounds.

As mouse lifespan experiments often have different results based on the sex of the animals tested ^33,34^, when required we created different instances for the same compound based on the sex of the mice used in the experiment. This greatly increases the number of examples available in the dataset and avoids conflating data that could lead the classification model to find incorrect patterns. However, the trade-off is that the feature values for all other features (except the sex feature) are the same among such instances, which can hinder the classifier’s performance if mixed-sex data is used to train the model.

Each instance in the datasets is assigned a binary class label indicating presence (class 1, positive) or absence (class 0, negative) of substantial evidence of lifespan increase when the compound was used in treatment during the longevity experiments reported in the papers. Naturally, the labelling process depends on many variables, as do the experiments performed in the source studies. The process we used to get a consistent class label definition for each instance, based on DrugAge entries, was as follows:

First, for each peer-reviewed publication studying the effect of a compound on mice, we analysed all reported results for each compound-sex combination. If at least one dosage or treatment regimen produces a median (or mean, where the median is not available) lifespan extension that is greater than 5% compared to the control mice population and is statistically significant (with p-value ≤ 0.05), then we count that as a positive result, otherwise we count it as a negative result.

Then, the results from different publications are aggregated so that each compound-sex combination is assigned a single class label, as follows. If the number of positive results is greater or equal to the number of negative results, the instance for that compound-sex combination gets a positive class label. Otherwise, it is labelled as the negative class (i.e., no substantial evidence of lifespan increase).

In summary, each instance in our dataset represents a combination of a compound and the sex of the mice used in the experiments (mixed-sex datasets have from 88 to 158 instances, depending on availability of data). An instance’s class label is defined by the consensus of the experimental results from DrugAge *M. musculus* section referring to that compound-sex combination, with positive labels indicating evidence of a positive association with longevity for treatments using that compound.

### 1.2 Dataset Preparation and Descriptions

To create our datasets, we created predictive features obtained from three source databases, namely STRING ^25^, LINCS ^26^ and PubChem ^27^. The features are different types of biological and chemical descriptors of compounds and their targets, all of which have been applied to drug discovery before. In this Section we detail the data collection processes for each of these feature types.

The STRING-based datasets use binary feature values from the descriptors (i.e., Gene Ontology, protein domains, biological pathways and keywords) of the targets of each compound, i.e., the proteins a compound was designed to interact with or known to be the main mediators of its pharmacological action. We obtained the list of a compound’s targets using several different sources, as detailed later in this Section. Additionally, the target descriptors are collected from STRING in two ways (functional annotation and enrichment), resulting in substantially different datasets.

The LINCS-based datasets have human gene expression data; consolidated numeric representations of over- and under-expression of each human gene from cell-line experiments after exposure to a compound. The consolidated values represent gene expression trends over all experiments with different configurations (cell-lines, doses and times).

Finally, the PubChem-based dataset is the only dataset of chemical descriptors used in our study. It has a set of binary variables, called Molecular Fingerprints, each one representing whether a compound’s structure contains a given chemical substructure or pattern.

For the datasets with binary feature values (i.e., the STRING-based datasets and Molecular Fingerprints) there are several features with too little variation in their values across the instances, which have little to no predictive value for a classification model. Therefore, as a data cleaning step prior to training our models, we applied a simple frequency-threshold filter to remove features with fewer than 3 instances with a “1” value (or fewer than 3 instances with a “0” value). After applying the frequency-threshold filter we also removed instances that had only “0” values in all remaining features, as they could no longer be distinguished by the classifiers.

#### 1.2.1 Datasets of STRING Descriptors of Protein Targets

We created a dataset based on target proteins associated with each compound and their descriptors, as follows. Firstly, instead of considering protein interactors listed on STITCH (a compound-protein interaction database ^35^, associated with STRING), which include predicted compound-protein interactions based on different levels of evidence, we have focused on identifying the specific targets of a compound, in order to create predictive features which are intuitively more biologically relevant. In order to cover the majority of compounds in DrugAge mice data and to obtain a reliable list of targets for each of them, we combined different sources for obtaining the list of targets of each compound:

- **DrugBank** ^28^: target data obtained from the DrugBank database’s website, accessed 01/2024;
- **Pharos** ^36^: target data obtained using the Pharos GraphQL API, Pharos v3.18.1;
- **BioGRID** ^37^: target data obtained through the database’s protein target table, v4.4.228;
- **ChEMBL** ^38^: target data obtained using the *chembl_webresource_client* python library ^39^;
- **Therapeutic Target Database** ^40^: target data obtained using the TTD database’s website, accessed 01/2024;
- **Source publications** for the compound’s DrugAge ^23^ entry and other relevant publications about the compound in case only one or no targets were identified via the sources listed above

Note that all target proteins obtained are for *Homo sapiens*, not *Mus musculus*, since the above databases focus on human data. After consolidating the target lists for all compounds, we used the STRING database API (v12.0) to obtain the protein descriptors for those targets through either functional annotation (FA) or neighbour enrichment (NE). In both FA and NE, STRING was queried for each protein target individually. However, unlike simple annotations returned by FA calls, NE calls returned statistically significant enrichments of annotations for the target protein and its 10 nearest functionally associated neighbours (automatically added by STRING).

The STRING API response collates annotations for the input proteins from various sources, related to their structure and function. We selected a subset of the feature types included in the API’s response, excluding those that were not relevant for our classification problem of drug discovery for longevity studies. The following feature types were used to create the binary predictive features in the Targets datasets:

- **Gene Ontology (GO) Terms**: These are widely used in similar drug discovery applications, as broad descriptors of genes’ structure and functions. We included all three categories of GO terms ^41^: Biological Processes, Molecular Functions and Cellular Components.
- **Protein Domains**: These features also represent information about the structure and function of proteins. The domains represented in the created dataset are from InterPro ^42^. Pfam and SMART (Simple Modular Architecture Research Tool) protein domains are also present in the STRING API responses, but these were not included, the former because they were incorporated by InterPro and the latter because there were too few annotations for them.
- **Pathways**: These features contain biological pathway information from Reactome ^43^, WikiPathways ^44^ and KEGG ^45^. Note that KEGG pathway features are not present in the FA datasets due to KEGG license restrictions keeping them from being returned in that call.
- **Uniprot Keywords**: These features represent a hierarchical set of keywords used to describe a protein entry on Uniprot ^46^.

In total we have 8 feature types in the Targets datasets, resulting in a considerably large number of features in each of them: 8,228 features in the FA dataset and 6,459 features in the NE dataset, after applying the minimum-threshold filter (these values refer to the full datasets, with 143 male and female mice instances). In order to assess the individual predictive performance of each feature type, we also did experiments with datasets prepared with single feature types.

#### 1.2.2 Datasets of LINCS Gene Expression

The Library of Integrated Network-Based Cellular Signatures (LINCS, v1.1) ^26^ dataset stores, among other types of data, *Homo sapiens* gene expression signatures. We created two datasets using the LINCS data, one including all genes (12,328 genes) and another using a subset of landmark genes called L_1000 (978 genes) ^47^. The latter refers to the genes for which the gene expression values were directly measured by the experiments, rather than inferred by the database designers.

For each gene expression experiment result stored in the LINCS database, the features are numeric values representing the (measured or estimated) expression of each human gene in cells belonging to a given cell line, after exposure to a dose of a given compound for a specified amount of time. Often dozens of experiments are performed for each compound, with different doses, times and cell lines. Therefore, a single compound can be associated with up to hundreds of entries on LINCS, with different gene expression values in each entry.

In order to use the gene expression values from LINCS in our machine learning context where each instance refers to a compound-sex combination, we need to have a single value for each gene (i.e., one row per compound), which broadly represents its gene expression after exposure to the compound. Hence, for each gene, we consolidated its expression value for each compound through a two-step process, as follows.

First, we selected the gene expression from the median dose over all entries in each cell line as the signature value for that cell line. Then, we calculated the median gene expression value over all cell lines’ signatures values. This is repeated for all genes in the dataset so that, after performing these steps, each gene has a single expression value for each compound. The numeric values vary roughly between -2 and +2, with positive values indicating an over-expression of the gene and vice-versa.

Naturally, this data consolidation process incurs some data loss through reducing the granularity of the LINCS data, and the validity of the aggregated values depend on many factors such as the standard deviation of the gene expression values measured over different experiments. On the other hand, the LINCS datasets have numerical values for their features, which may be advantageous for decision tree-based algorithms (like the algorithm used in this work) because the algorithm has more options for partitioning the data based on a feature’s values.

#### 1.2.3 Dataset of PubChem Molecular Fingerprints

Studies that apply machine learning for drug discovery problems often use chemical substructures of compounds as predictive features, either by themselves or in combination with biological data ^8–10^. In our study, we created datasets where the binary features represent the absence/presence of 880 chemical substructures in a given compound. Importantly, the feature values for chemical data are independent from the model organism and, in the case of the substructures used in our study, easily obtainable (Molecular Fingerprints are available for any compound present in PubChem).

The Molecular Fingerprints substructures may represent an element count (e.g., compound has ≥ 8 hydrogen atoms), a type of ring system (e.g., compound has saturated or aromatic carbon-only ring size 3), atom pairings, neighbourhoods and bonding (e.g., compound has Si – Cl bond), and other such patterns. The data was obtained from the PubChem Substructure Fingerprint data, using the *pubchempy* python library (v1.0.4).

### 1.3 Experimental Setup for Machine Learning

We trained Random Forest (RF) classifiers ^48^ using each of the datasets produced for this study, using the same experimental setup described in this Section. Ensembles of classifiers based on decision trees (like RFs) often outperform more complex algorithms such as deep learning when applied to tabular data for binary classification ^49^, and RFs also have the advantage of being much faster than deep learning algorithms in general. In addition, although RFs are not as interpretable as single decision trees, they still retain some degree of interpretability, mainly through feature importance metrics that take into account the contents of the trees in the forest (i.e. the model’s internal mechanisms).

For all experiments we performed a 5-fold cross-validation process for each RF classifier and ran 10 different experiments for each dataset (varying the random seed parameter, resulting in different training/test data divisions and different RFs). The rationale for repeated experiments was to obtain a more robust estimation of the models’ predictive accuracy, as our datasets have few instances and, therefore, there is more stochastic variation on each models’ predictive accuracy estimation.

In Section 3 we report the average values over all folds (5 folds over 10 cross-validation runs) of two predictive accuracy metrics: the Area Under the Receiver Operating Characteristic Curve (AUC) ^50^ and the Geometric Mean of Sensitivity and Specificity (G-mean). Both metrics vary between 0 and 1, with 1 representing a perfect classifier. The AUC is widely used as a general representation of a binary classification model’s performance, and it has the advantage of providing a measurement that is independent of the threshold used for the class-label decision. When used by itself the AUC may result in over-optimistic evaluations of models that favour the majority class, so we also used the G-mean as a conservative metric that assigns equal weight to both class labels and considered both metrics when evaluating our RF models.

The RFs were trained using the *sklearn* implementation (v1.4.0) ^51^, with 500 random trees and the default values of all other hyperparameters. The datasets created for this research have a class-imbalance issue (between 22% and 47% positive-class instances, and so between 78% and 53% negative-class instances, depending on the dataset). Although none of the imbalances are very strong, this can still lead to models biased in favour of the majority class (the negative class, representing compounds without a known association with longevity). Therefore, in each RF, we increased the weight of the minority class instances in the training set until a balance was reached, meaning the classifier penalized misclassifications of minority class instances more severely. This was preferable to the common alternative method of undersampling majority class instances, as class-weight adjustment does not reduce the number of examples in the dataset.

After training and evaluating our RF models, we selected a group of 10 models (5 from mixed-sex datasets and 5 from male-only datasets) that had the best predictive performance, to analyse how they made their predictions and to identify compounds with potential for lifespan extension in mice.

First, we ranked all features used for each of the selected models, based on each feature’s ability to discriminate between the positive-label and negative-label examples, by using the default feature importance measure in sklearn (the average class-impurity decrease measure ^52^). Then, the top features in that ranking are identified as the most important features for that model.

Next, we used the selected models to identify promising compounds for future studies of mice longevity, in two ways. The first was identifying the consistent false-positive classifications (i.e., over half of the top models classified the compound as positive whilst the compound is annotated in the data as negative) in the labelled DrugAge data used in these experiments. The second way was using the selected models as ensembles of RFs to classify unlabelled data from an external dataset and identifying the most confident positive-class classifications by these ensembles. In order to discuss the over 200 compounds identified in the external dataset analysis, we used the UMAP (Uniform Manifold Approximation and Projection) dimension reduction technique ^53^ to represent each data point (compound) in a two-dimensional space based on their similarities. Then, we grouped these data points using the DBSCAN (Density-Based Spatial Clustering of Applications with Noise) clustering algorithm ^54^ to find cohesive clusters of similar compounds. Each of these analyses is discussed in Section 3.

### 1.4 Testing predictions in *C. elegans*

Compounds were purchased in powder form from MedChemExpress LLC, USA. The compounds ordered were: BMS-754807 (HY-10200), Umeclidinium bromide (HY-12100), Sumanirole maleate (HY-70081A), Tolterodine (HY-A0024), Rotigotine (HY-75502), Voxtalisib (HY-15900), Pramipexole (HY-B0410), Dofetilide (HY-B0232), Solifenacin (HY-A0034), Brinzolamide (HY-B0588), Epinastine (HY-B0640), Bromocriptine mesylate (HY-12705A), Cabergoline (HY-15296), Aclidinium bromide (HY-14144), Quinagolide hydrochloride (HY-13736A), Tiotropium bromide (HY-17360), Glycopyrrolate (HY-17465), Dorzolamide hydrochloride (HY-B0109A), Darifenacin hydrobromide (HY-A0012), Sarizotan (HY-100820). Mianserin (M2525) and Dihydroergotamine Mesylate (1202005) were purchased in powder form from Millipore Sigma, USA. Age-synchronized *C. elegans* were cultured in liquid medium and dispensed into flat-bottom, optically clear 96-well plates (Corning 351172) at ∼8 animals per well in 150 µL of 6 mg/mL X-ray-irradiated OP50. Animals were seeded as L1 larvae and grown at 20 °C with plates sealed to prevent evaporation. To block self-fertilization, FUDR (120 µM final; Sigma F0503) was added 42–45 hr after seeding. Drugs were added on day 1 of adulthood, with DMSO maintained at 0.5% (vol/vol). Survival statistics were calculated using the Mantel–Haenszel log-rank test.

## Results and Discussion

### 1.5 Predictive Accuracy Results

The goal of our study was to train Random Forest classifiers on the murine lifespan data from DrugAge so that they can be employed to classify compounds previously not tested in mice into potentially lifespan-extending or not. To this aim, we generated male-only, female-only and mixed-sex datasets, as there are substantial sex-related differences in lifespan responses to various compounds.

The average AUC and average G-mean results from 10 runs of 5-fold cross-validation experiments for models trained on each full mixed-sex or male-only dataset are presented in Table 1. Table 2 shows the average AUC and G-mean values from models trained on each individual Targets feature category (Functional Annotation, FA, or Neighbour Enrichment, NE, as defined in Section 2.2.1).

**Table 1.** Average AUC and G-mean values from models trained on each full dataset.

| Dataset | Mixed-sex |  | Male-only |  |
| --- | --- | --- | --- | --- |
|  | AUC | G-mean | AUC | G-mean |
| Targets FA (Functional Annotation STRING descriptors) | 0.678 | 0.327 | 0.626 | 0.472 |
| Targets NE (Neighbour Enrichment STRING descriptors) | 0.636 | 0.392 | **0.63** | **0.6** |
| LINCS gene expression (all genes) | 0.564 | 0.135 | 0.557 | 0.469 |
| LINCS gene expression (L_1000 landmark genes) | 0.634 | 0.161 | 0.568 | 0.518 |
| PubChem Molecular Fingerprints | **0.637** | **0.472** | 0.636 | 0.497 |

**Table 2.** Average AUC and G-mean values from models trained on each individual feature category.

| Dataset | Mixed-sex |  |  |  | Male-only |  |  |  |
| --- | --- | --- | --- | --- | --- | --- | --- | --- |
| Feature preparation | Functional Annotation |  | Neighbour Enrichment |  | Functional Annotation |  | Neighbour Enrichment |  |
| Performance metric | AUC | G-mean | AUC | G-mean | AUC | G-mean | AUC | G-mean |
| GO Terms – Biological Process | 0.692 | 0.317 | 0.621 | 0.37 | 0.657 | 0.498 | **0.634** | **0.594** |
| GO Terms – Molecular Function | 0.6 | 0.334 | 0.618 | 0.338 | 0.548 | 0.471 | 0.636 | 0.498 |
| GO Terms – Cellular Component | 0.629 | 0.364 | **0.653** | **0.474** | 0.571 | 0.465 | **0.657** | **0.528** |
| Uniprot Keywords | 0.614 | 0.306 | 0.598 | 0.37 | 0.578 | 0.43 | 0.597 | 0.475 |
| InterPro Protein Domains | 0.589 | 0.483 | 0.555 | 0.468 | **0.663** | **0.534** | 0.559 | 0.488 |
| Wiki Pathways | 0.662 | 0.364 | **0.653** | **0.446** | 0.565 | 0.38 | 0.617 | 0.55 |
| Reactome Pathways | 0.601 | 0.354 | **0.648** | **0.522** | 0.548 | 0.43 | 0.627 | 0.577 |
| KEGG Pathways | N /A | N /A | **0.629** | **0.451** | N /A | N /A | **0.647** | **0.615** |

Note that we also ran experiments with female-only datasets, but the models resulting from these experiments were not valid (i.e., they were overfitted to the majority class, and thus unable to make reliable predictions for minority-class compounds), thus we are not reporting these results here. We believe that we did not have sufficient female-only instances of the positive class to train models that are able to make meaningful predictions for positive-class compounds – notable only around 20% of the female instances belonged to the positive class, whilst around 40% of the male instances belonged to the positive class.

In order to select models to be analysed in depth in this Section, we first set a minimum threshold of G-mean, as this metric is a good way to determine whether a model has been overfitted towards the majority-class. Using a secondary criterion to disregard overfitted results was required because some models achieved relatively high AUC values (e.g., 0.692 for FA Biological Process) but tended to skew predictions towards the majority (negative) class, which made their AUC results overoptimistic, and this is reflected in their low G-mean results (in this case, 0.317).

For the mixed-sex datasets, which had models with lower predictive accuracy overall, the minimum G-mean threshold was set to 0.4. For the male-only datasets we set it to 0.5, as these models performed better overall. Then, we selected the best 5 models that passed the G-mean threshold from each group of experiments (mixed-sex datasets and male-only datasets), based on their AUC results. The 10 selected models have their results highlighted in boldface in Tables 1 and 2.

Overall, the top models have similar average AUC (between 0.629 and 0.663) values. However, their G-mean results show that the male-only models are stronger than the mixed-sex models, likely because the female mice instances are harder to classify correctly due to the lack of female positive-class instances, making the problem more difficult in mixed-sex data (as mentioned earlier, ∼20% of the female-mice instances in our data are positive, compared to ∼40% of the male-mice instances). The similar AUC values between models that have such different G-means can be explained by a tendency of AUC to reward correct classifications of the majority class, so conservative models that tend to classify instances as negative (a safer classification, as negative instances represent a larger proportion of the data) are achieving good AUC results. An example of this issue is the GO Terms – Biological Process FA model in the mixed-sex data, which had the highest AUC value overall (0.692) but a very low G-mean (0.317). Such models overfit to the majority class, and this is reflected by metrics such as the G-mean, which is why we consider both AUC and G-mean when evaluating our models.

The LINCS datasets did not yield good models in any of the experiments, which we believe is due to the higher complexity of the data (numerical values of gene expression rather than binary values), which makes the small number of available examples take a more significant toll on the classifier’s performance. Another explanation could be that gene expression averaged across many cell lines, dosages and timepoints is a poor representation of longevity effects of tested compounds.

In the mixed-sex datasets, the Molecular Fingerprints model (based on chemical substructures in the compounds’ compositions) was selected as one of the 5 best models, whilst all other selected models belong to the Targets category (based on STRING data of protein target annotations).

In the Targets datasets, the NE mode of dataset creation, which includes annotations (GO Terms, protein domains, pathways and keywords) common to each target’s neighbourhood, achieved better results than the FA mode, which includes only the annotations from the input targets themselves. 8 out of 9 top models in Targets category used the NE mode, with the exception of the FA InterPro dataset in male-only model. The NE features refer to annotations that are common to a group of proteins, which makes them more selective compared to FA annotations. This may be the reason for this disparity between the success of NE and FA models. It is also worth noting that the selected male-only models included the NE dataset combining all 8 feature categories (GO annotations, biological pathways, protein domains and UniProt keywords), with a relatively high average G-mean of 0.6, the second-best G-mean result over all experiments. This shows that even feature categories that did not generate a top model on their own had value when considered in combination with other categories. Moreover, GO Function and Uniprot Keywords have not been selected for any of the top models, likely indicating their low informativeness for the task of predicting longevity drugs.

### 1.6 Analysis of Feature Importance in the Best Predictive Models

Tables 3–7 show, for each of our 5 selected models trained on male-only datasets, which 10 features had the highest importance when labelling a compound as positive (predicted to increase lifespan in mice) or negative class. Please note that as only top 10 features are listed for each model, the lists of features discussed are not comprehensive or exhaustive. We present the results of models trained on mixed-sex datasets in the Supplementary information due to space limitations.

**Table 3.** Most relevant features in the FA InterPro Domains model.

| Feature | Description | Class Tendency |
| --- | --- | --- |
| IPR017452 | GPCR, rhodopsin-like, 7TM | Positive class |
| IPR000276 | G protein-coupled receptor, rhodopsin-like | Positive class |
| IPR036291 | NAD(P)-binding domain superfamily | Negative class |
| IPR003193 | ADP-ribosyl cyclase (CD38/157) | Unclear |
| IPR002233 | Adrenoceptor family | Positive class |
| IPR016024 | Armadillo-type fold | Positive class |
| IPR014729 | Rossmann-like alpha/beta/alpha sandwich fold | Negative class |
| IPR029071 | Ubiquitin-like domain superfamily | Positive class |
| IPR018490 | Cyclic nucleotide-binding domain superfamily | Positive class |
| IPR000595 | Cyclic nucleotide-binding domain | Positive class |

**Table 4.** Most relevant features in the NE Gene Ontology Process model.

| Feature | Description | Class Tendency |
| --- | --- | --- |
| GO:0003008 | System process | Positive class |
| GO:0007191 | Adenylate cyclase-activating dopamine receptor signalling pathway | Positive class |
| GO:0042391 | Regulation of membrane potential | Positive class |
| GO:0019752 | Carboxylic acid metabolic process | Negative class |
| GO:0007189 | Adenylate cyclase-activating G protein-coupled receptor signalling pathway | Positive class |
| GO:0007186 | G protein-coupled receptor signalling pathway | Positive class |
| GO:0001503 | Ossification | Positive class |
| GO:0032879 | Regulation of localization | Positive class |
| GO:0043648 | Dicarboxylic acid metabolic process | Negative class |
| GO:0007212 | Dopamine receptor signalling pathway | Positive class |

**Table 5.** Most relevant features in the NE Gene Ontology Components model.

| Feature | Description | Class Tendency |
| --- | --- | --- |
| GO:0005834 | Heterotrimeric G-protein complex | Positive class |
| GO:0005886 | Plasma membrane | Positive class |
| GO:0045177 | Apical part of cell | Positive class |
| GO:0005829 | Cytosol | Negative class |
| GO:0016323 | Basolateral plasma membrane | Unclear |
| GO:0005739 | Mitochondrion | Negative class |
| GO:0005667 | Transcription regulator complex | Negative class |
| GO:1904813 | ficolin-1-rich granule lumen | Negative class |
| GO:0005759 | Mitochondrial matrix | Negative class |
| GO:0090575 | RNA polymerase II transcription regulator complex | Unclear |

**Table 6.** Most relevant features in the NE KEGG Pathways model.

| Feature | Description | Class Tendency |
| --- | --- | --- |
| hsa04929 | GnRH secretion | Positive class |
| hsa04911 | Insulin secretion | Positive class |
| hsa04022 | cGMP-PKG signalling pathway | Positive class |
| hsa05146 | Amoebiasis | Unclear |
| hsa05152 | Tuberculosis | Negative class |
| hsa05032 | Morphine addiction | Positive class |
| hsa04742 | Taste transduction | Positive class |
| hsa04914 | Progesterone-mediated oocyte maturation | Positive class |
| hsa04713 | Circadian entrainment | Positive class |
| hsa04640 | Hematopoietic cell lineage | Negative class |

**Table 7.**
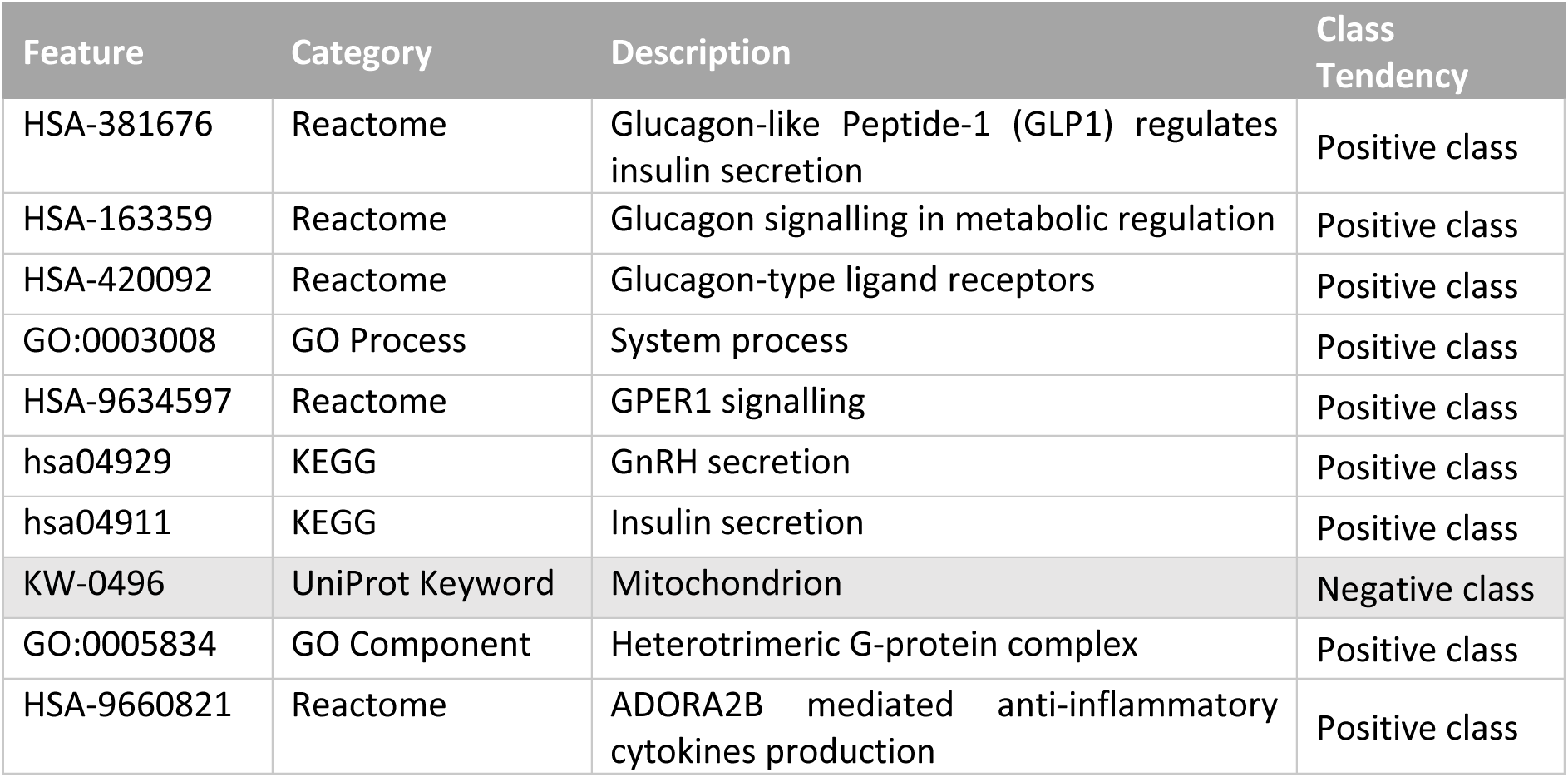
Most relevant features in the NE All categories model.

| Feature | Category | Description | Class Tendency |
| --- | --- | --- | --- |
| HSA-381676 | Reactome | Glucagon-like Peptide-1 (GLP1) regulates insulin secretion | Positive class |
| HSA-163359 | Reactome | Glucagon signalling in metabolic regulation | Positive class |
| HSA-420092 | Reactome | Glucagon-type ligand receptors | Positive class |
| GO:0003008 | GO Process | System process | Positive class |
| HSA-9634597 | Reactome | GPER1 signalling | Positive class |
| hsa04929 | KEGG | GnRH secretion | Positive class |
| hsa04911 | KEGG | Insulin secretion | Positive class |
| KW-0496 | UniProt Keyword | Mitochondrion | Negative class |
| GO:0005834 | GO Component | Heterotrimeric G-protein complex | Positive class |
| HSA-9660821 | Reactome | ADORA2B mediated anti-inflammatory cytokines production | Positive class |

In all tables we included a Class Tendency column indicating whether features clearly increased the chance of a certain class label. The class tendency of each binary feature F was calculated using two metrics we called Effectiveness (ratio of instances of the positive class (*C*=1) among instances that have the feature present (*F*=1)) and Dispensability (ratio of instances of the positive class among instances that have the feature absent (*F*=0)). The result of the division between those metrics indicates an increase/decrease of likelihood of a positive-class classification when using the feature’s value to split the data. We set cut-off values to determine whether the feature’s value results in a tendency towards each class, as shown in Equation 1.

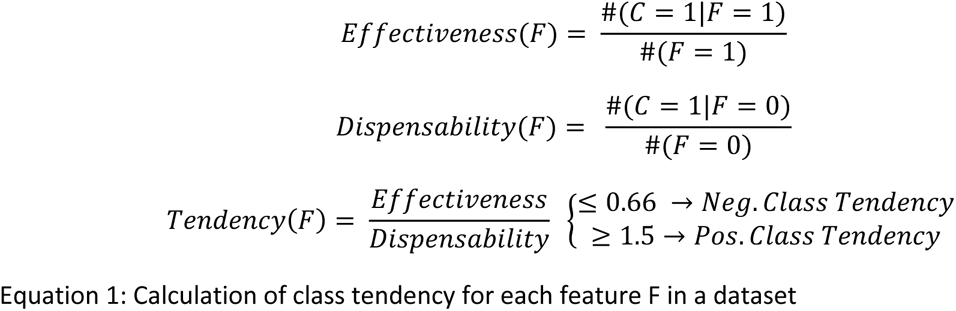

As an example of this tendency calculation, consider the ‘IPR017452’ feature in the FA InterPro Domains model (top-ranked feature in Table 3): out of the 21 instances with value ‘1’ for this feature, 13 are of the positive class (Effectiveness = 13/21 = 61.90%); out of the 49 instances with value ‘0’ for this feature, 12 are of the positive class (Dispensability = 12/49 = 24.49%). Therefore, the class tendency of ‘IPR017452’ is 2.53 (61.90/24.49), which indicates a positive-class tendency for this feature.

Table 3 demonstrates that compounds interacting with rhodopsin-like G protein-coupled receptors (GPCRs), including adrenoceptors, as well as proteins containing Armadillo-type folds, ubiquitin-like domains, or cyclic nucleotide-binding domains, are promising for lifespan extension. GPCRs are essential in signal transduction, influencing pathways related to metabolism, stress response, and hormonal regulation, all of which are vital for maintaining homeostasis. It has to be noted that GPCRs are very common drug targets in general, so their enrichment here could in part be attributed to their excellent pharmacological properties. Still, they had positive class tendency, meaning they were more likely to be found amongst drugs extending lifespan rather than amongst drugs with null or negative effects on lifespan. Armadillo-type fold proteins, such as those involved in the Wnt signalling pathway, contribute to cellular proliferation, differentiation, and maintaining tissue homeostasis through their role in cell adhesion and cytoskeletal integrity. This is consistent with the mixed-sex NE Wiki Pathways model, where “Wnt signalling pathway and pluripotency” was identified as a top predictive feature (Supplementary Table 2). Ubiquitin-like domains are crucial for proteostasis, aiding in protein quality control and cellular stress responses. Cyclic nucleotide-binding domains, involved in binding cAMP and cGMP, regulate metabolic processes and enhance cellular resistance to oxidative stress. This is consistent with the mixed-sex NE KEGG Pathways model where “cGMP-PKG signalling pathway” was identified as a top predictive feature (Supplementary Table 4). Compounds targeting proteins with NAD(P)-binding domains or Rossmann-like alpha/beta/alpha sandwich folds are less promising for lifespan extension. These domains are primarily involved in redox reactions and basic metabolic processes, which, while essential for cellular function and important to longevity, do not seem to make good drug targets to modulate longevity.

The top feature in Table 4 is the very broad “System process” GO term, which indicates that compounds which target multicellular organismal processes carried out by organs or tissues within an organ system, such as immune, circulatory, nervous, etc., have the most potential to extend lifespan. Specific processes worth targeting for lifespan extension include G protein-coupled receptor signalling pathways, specifically adenylate cyclase-activating ones, in particular the dopamine receptor pathway, as well as regulation of membrane potential, ossification and “regulation of localization”. The “Regulation of membrane potential” feature is consistent with the “Plasma membrane” feature from the NE Gene Ontology Components models (Table 5, Supplementary Table 1) and the “Potassium channels” feature from the mixed-sex NE Reactome Pathways model (Supplementary Table 3). “Regulation of localization” is a GO term that describes “any process that modulates the frequency, rate or extent of any process in which a cell, a substance, or a cellular entity is transported to, or maintained in, a specific location”. Targeting carboxylic and dicarboxylic acid metabolism in a manner that extends lifespan is likely to be difficult as different tissues have different metabolic and mitochondrial demands that may be hard to modulate without causing negative effects in at least one tissue These chemicals are important intermediates in various metabolic pathways, including the Krebs cycle, which is essential for ATP production, cellular respiration and overall metabolic health. This is consistent with NE Gene Ontology Components models (Table 5, Supplementary Table 1), where the mitochondrial matrix (where the Krebs cycle occurs) was identified as a negative class feature.

Table 5 shows results from the NE Gene Ontology Components model applied to a male-only dataset and is highly similar to Supplementary Table 1 which shows results from the same model but applied to a mixed-sex dataset. According to these results, compounds interacting with proteins localised at the plasma membrane and/or apical part of the cell, especially the heterotrimeric G-protein complexes, are likely to extend lifespan. G-protein complexes are essential signalling molecules functioning as molecular switches to transmit signals from cell surface receptors (which are thus called G protein-coupled receptors) to various intracellular effectors. They are involved in numerous physiological processes, including sensory perception, immune responses, and regulation of mood and metabolism. Interestingly, *GNAQ, GNA11* and *GNAS* – genes that encode different alpha subunits of heterotrimeric G-protein complexes – have recently been predicted to be some of the strongest cancer drivers ^55^, so inhibiting them with chemical compounds might extend murine lifespan by delaying cancer development. Compounds interacting with proteins localised in the cytosol, mitochondrion, especially mitochondrial matrix, ficolin-1-rich granule lumen, or transcription regulator complex are underrepresented either because these proteins have no role in longevity or because targeting them poses pharmacological difficulties.

Table 6 shows results from the NE KEGG Pathways model applied to a male-only dataset and is similar to Supplementary Table 4 which shows results from the same model but applied to a mixed-sex dataset. According to these results, compounds interacting with proteins involved in gonadotropin-releasing hormone (GnRH) secretion, progesterone-mediated oocyte maturation, insulin secretion, cGMP-PKG signalling pathway, taste transduction, circadian entrainment and morphine addiction are promising candidate lifespan-extending compounds. It is peculiar that the mixed-sex NE Wiki Pathways model classified “Circadian rhythm genes” as a negative class feature (Supplementary Table 2), which seems to be at odds with the current model (Table 6) classifying “Circadian entrainment” as a positive class feature. Morphine addiction was selected likely because it is mediated by dopamine, and thus this result is in agreement with the “Dopamine receptor signalling pathway” feature from the NE Gene Ontology Process model (Table 4). As taste transduction can affect the palatability of food, it may indicate that some longevity drugs in our training dataset affect the feeding behaviour of mice and thus cause indirect (behavioural) caloric restriction, which is known to strongly extend lifespan. For example, it has been shown that dietary restriction response and lifespan extension can be achieved through the pharmacological masking of a sensory pathway that signals the presence of food ^56^. Sensory perception in general has been shown to modulate lifespan in worms^57^ and flies^58^. Compounds targeting proteins involved in tuberculosis and hematopoietic cell lineage are unlikely to be effective for lifespan extension.

Finally, the NE All categories model (Table 7) was able to select features from all categories, and thus most of them have already appeared as top features in single-category models. We can see that these features are mostly related to systemic processes, such as hormone secretion and signalling mediated via G-protein coupled receptors. The top hormones involved are Glucagon-like Peptide-1 (GLP1), glucagon, insulin, oestrogen (signalling via G protein-coupled estrogen receptor 1 (GPER1)), gonadotropin-releasing hormone (GnRH) and adenosine (signalling via adenosine A2B receptor (ADORA2B)). Notably, all of them signal via G protein-coupled receptors. These results lend support to hormone theories of ageing ^59,60^. As three of these hormones are involved in glucose metabolism, this is consistent with the occurrence and detrimental effects of diabetes and metabolic syndrome with age, as well as the effectiveness of caloric restriction and antidiabetic drugs such as acarbose and possibly metformin. Two other hormones are sex hormones, which aligns with sex-specific differences observed in human and animal lifespan and in the effectiveness of lifespan-extending compounds. The natural ligand for adenosine receptor A2B (ADORA2B) is extracellular adenosine, formed from the reduction of ATP by ENTDPases. ATP enters the extracellular space in response to tissue injury and apoptosis amongst other stress factors and has chemotactic and excitatory effects. The reduction of ATP to adenosine is thought to be a regulatory mechanism by which the synthesis of anti-inflammatory cytokines is induced ^61^. Thus, the “ADORA2B mediated anti-inflammatory cytokines production” feature, together with the “Chemokine signalling pathway” feature from mixed-sex NE Wiki Pathways (Supplementary Table 2) and NE KEGG Pathways (Supplementary Table 4) models, lend support to the inflammageing theory and highlights the importance of reducing chronic inflammation^62^. Although the UniProt Keywords model did not perform as well as other models that we selected, the identification of the UniProt Keyword “Mitochondrion” feature as a negative class one in the current combined model (Table 7) is consistent with the results of the NE Gene Ontology Components models (Table 5, Supplementary Table 1), which also nominated mitochondrion and mitochondrial matrix as negative class features. These results suggest a critical look is required at the mitochondrial theory of ageing.

### 1.7 Identifying the Most Promising Novel Compounds for Lifespan Extension

After identifying the models with the best predictive accuracy in Section 3.1, in this Section we use their predictions to highlight compounds that show some promise of lifespan-extension capabilities.

A classifier predicts an instance as part of the positive class based on feature patterns that indicate its similarity to training instances of that class, so the compounds identified in this analysis have some commonalities with previously successful compounds. Naturally, this does not guarantee that the compounds would work similarly. For example, they may interact differently with the target (e.g., activate it whilst the successful instances inhibit it), and this would not be reflected in our binary-feature datasets.

#### 1.7.1 Identifying recurrent false-positive classifications in top models

In order to identify novel compounds that may extend murine lifespan, we first considered the false-positive classifications in our own models’ predictions, as these indicate that the model identified positive-class patterns in the compound’s data. Table 8 shows six compounds that were frequently classified as longevity-related but are part of the negative class in our datasets (i.e., there is currently no sufficient evidence to claim that these compounds extend the median lifespan of mice). It includes the PubChem ID of each compound, and the ratio of the number of times the compound was classified as a false-positive over the number of top-model datasets it was included in (Putrescine and Chlorpheniramine are included only in mixed-sex datasets, as they only have examples from female mice studies).

**Table 8.**
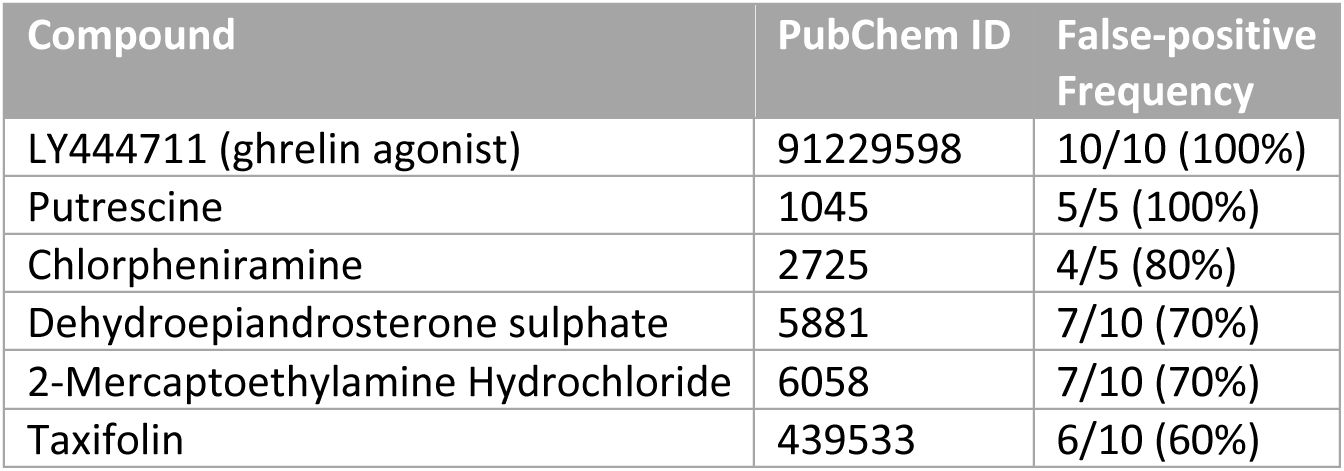
Negative-class compounds consistently classified as false-positive by the selected models.

These compounds have been reported as unsuccessful for lifespan extension in mice (see our positive-class definition in Section 2.1), but they are consistently being labelled as positive by our most accurate models, which shows they have some similarity with positive-class examples and may warrant further investigation. Note that although the negative-class compounds have failed to extend lifespan in the existing studies, it is possible that a different treatment regimen (e.g. a different dosage, route of administration, starting age, duration) or a different mouse strain could lead to positive results. Alternatively, these drugs might have some off-target effects that negatively impact lifespan. Interestingly, LY444711 slightly improves median survival in mice, although the increase does not reach statistical significance. What is remarkable is that it significantly improves maximal lifespan ^63^, although we do not use that parameter for training the models. Another drug in our false-positive list, dehydroepiandrosterone sulphate, has been shown to be ineffective in two mouse studies, but its levels are dramatically reduced with age in humans ^64^ (perhaps it is one of the strongest blood biomarkers of ageing), so it is still possible that the right concentration or treatment regimen has not been achieved, given that this is a hormone with complex and precisely regulated secretion patterns^65^. Moreover, as rodents secrete small amounts of DHEA, it has been suggested they cannot serve as a research model for this hormone ^59^. It could also be that DHEA downregulation with age is a protective response that is activated by the organism to fight aging, so supplementing it might have negative consequences.

#### 1.7.2 Labelling unseen data from an external dataset

For this analysis, we created two classifier ensembles using the selected (most accurate) classification models from each type of dataset: 5 from mixed-sex datasets and 5 from male-only datasets. Each model outputs a positive-class prediction probability between 0 and 1, and the ensemble’s final probability is the average over all valid models’ probabilities (i.e., in cases where a model has no data for a compound, it is not included in the ensemble’s result calculation).

We used these ensembles to classify a large number of existing drugs and compounds that have never been tested for their lifespan extension effects in mice. We downloaded the full DrugBank ^28^ dataset of Drug Target Identifiers (version 5.1.12, released on 2024-03-14) and used it to create feature datasets from STRING annotations and enrichments for these targets. After applying our ensembles to these datasets, we selected all novel compounds that were predicted by the ensembles to have at least 75% positive-class probability, which resulted in a list of 57 compounds from the mixed-sex ensemble and 272 compounds from the male-only ensemble. The male-only ensemble predicted more high-confidence compounds likely because those models had higher discriminatory power (G-mean). Some compounds confidently classified as belonging to the positive class by these ensembles are very similar to previously successful compounds (i.e., sharing targets with positive-class instances in the training data), whereas other compounds might represent previously unexplored research directions for future longevity studies in mice. Compounds with 100% feature similarity to the ones in the training dataset were removed.

As mentioned in Section 2.3, we used the UMAP dimensionality reduction technique to visualise the selected compounds in a two-dimensional space based on measuring Jaccard distances (a similarity metric for binary data that disregards ‘0’ matches) between the compounds. We adjusted the UMAP parameters to emphasise the global structure as opposed to local structure (i.e. set neg_sample_rate and n_neighhbours equal to the number of compounds minus one). Then, we used the DBSCAN algorithm to identify clusters of compounds, based on their Euclidean distance within the UMAP projection. We fine-tuned the DBSCAN eps parameter to keep the lowest possible number of groupings while preserving the distinct sets of top targets in each cluster, resulting in 4 clusters out of the 57 compounds nominated by the mixed-sex ensemble and 7 clusters out of the 272 compounds selected by the male-only ensemble.

Figure 2 shows the UMAP projections of the most confident positive-class predictions of the ensemble trained on male-only datasets, as well as the positive class likelihoods of these compounds and their clusters as determined by DBSCAN. Table 9 shows the clusters of compounds with positive-class likelihood ≥75% as estimated by the ensemble trained on male-only datasets, and each cluster’s most frequent protein targets are shown in Table 10. Due to space limitations, we present results for the ensemble trained on mixed-sex datasets in the Supplementary information.

**Figure 2.**
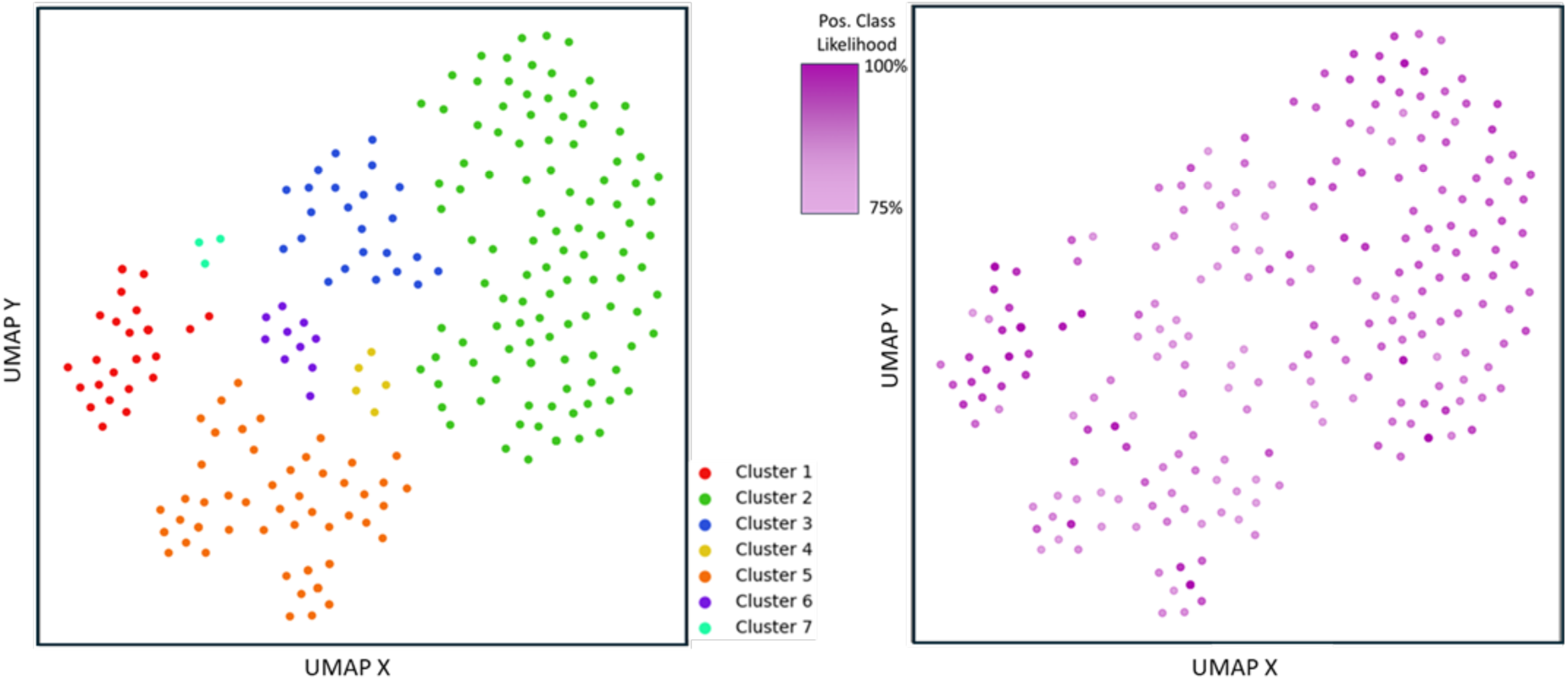
Clustering (left) and positive-class likelihood (right) of DrugBank compounds with positive-class likelihood ≥75% predicted by the ensemble trained on male-only datasets

**Table 9.** Clusters of DrugBank compounds with positive-class likelihood ≥75% nominated by the ensemble trained on male-only datasets. Potential lifespan-extending compounds are highlighted in bold.

| Cluster | Number of compounds | Positive class likelihood Min-Max (Mean) | Compounds |
| --- | --- | --- | --- |
| 1 | 37 | 75.3% to 97.2% (mean: 90.6%) | 2-hydroxymethyl-6-octylsulfanyl-tetrahydro-pyran-3,4,5-triol, **Acebutolol**, **Alprenolol**, **Arotinolol**, **Atenolol**, **Befunolol**, **Betaxolol**, **Bethanidine**, **Bevantolol**, **Bisoprolol**, **Bopindolol**, **Carteolol**, **Celiprolol**, **Cryptenamine**, Dipivefrin, DL-Methylephedrine, Droxidopa, Ephedrine, Isoetharine, Isoprenaline, **Labetalol**, **Levobunolol**, Mephentermine, **Metipranolol**, Mirabegron, **Nadolol**, Norepinephrine, **Penbutolol**, Phenoxybenzamine, Phenylpropanolamine, **Pindolol**, Pirbuterol, **Propafenone**, **Propranolol**, Racepinephrine, **Sotalol**, **Timolol** |
| 2 | 119 | 76.2% to 91.2% (mean: 83.9%) | Acepromazine, Alizapride, **Amantadine**, Amisulpride, Amitifadine, Amitriptyline, Amoxapine, Amphetamine, Aniracetam, **Apomorphine**, Aripiprazole, AS-8112, Asenapine, Bicifadine, Bifeprunox, BL-1020, Blonanserin, Brasofensine, Brexpiprazole, **Bromocriptine**, Bromopride, Buspirone, **Cabergoline**, Cariprazine, Carphenazine, Chlorprothixene, Cinnarizine, Clozapine, Desipramine, **Dihydro-alpha-ergocryptine**, Dihydroergocornine, Dihydroergocristine, **Dihydroergotamine**, Domperidone, Dopamine, Doxepin, Droperidol, Ephedra sinica root, Ergoloid mesylate, Ergotamine, Escitalopram, Etoperidone, Fluoxetine, Flupentixol, Fluspirilene, Haloperidol, Hydrocodone, Iloperidone, Imipramine, JNJ-37822681, Ketamine, Ketobemidone, **Lisuride**, Loxapine, Lumateperone, Lurasidone, Maprotiline, Melperone, Memantine, Meperidine, Mesoridazine, Methadone, Methotrimeprazine, Metoclopramide, **Mianserin**, Minaprine, Molindone, Naltrexone, Nefazodone, Norclozapine, Nortriptyline, Ocaperidone, Olanzapine, Ondansetron, Orphenadrine, Paliperidone, Pardoprunox, Paroxetine, Pergolide, Perospirone, Pipamperone, Pipotiazine, **Piribedil**, **Pramipexole**, Prochlorperazine, Promazine, Propiomazine, Quetiapine, **Quinagolide**, Raclopride, Remoxipride, Risperidone, Rolicyclidine, **Ropinirole**, **Rotigotine**, **Sarizotan**, Sertindole, Setiptiline, Sulpiride, **Sumanirole**, Tetrahydrocannabinavarin, Tetrahydropalmatine, Thiethylperazine, Thioproperazine, Thioridazine, Thiothixene, Tianeptine, Tramadol, Triflupromazine, Trimipramine, Viloxazine, Vortioxetine, YKP-1358, Yohimbine, Ziprasidone, Zotepine, Zuclopentixol |
| 3 | 24 | 75.1% to 87.2% (mean: 80.1%) | Agmatine, **Azimilide**, **Bepridil**, Cocaine, **Dofetilide**, **Enflurane**, Flunarizine, Gabapentin, **Halothane**, Isavuconazole, **Isoflurane**, Lamotrigine, Loperamide, Nicardipine, Pentoxifyverine, Perphenazine, Pimozide, Promethazine, Ranolazine, Ritodrine, **Topiramate**, Trifluoperazine, Trimebutine, Verapamil |
| 4 | 5 | 75% to 78.5% (mean: 76.6%) | **4-Methoxyamphetamine**, **Benzatropine**, Benzphetamine, **Diphenylpyraline**, Metamphetamine |
| 5 | 74 | 75% to 89.3% (mean: 78.3%) | **Aclidinium**, ALKS 27, Apraclonidine, Aranidipine, Astemizole, **Atropine**, Benzquinamide, Bethahistine, Bethanechol, Brompheniramine, Butriptyline, Captodiame, Cinitapride, Cirazoline, Cisapride, Clonidine, Cyproheptadine, **Darifenacin**, Disopyramide, Dosulepin, Dotarizine, Doxazosin, Epicept NP-1, **Epinastine**, Esmirtazapine, Fenfluramine, Fenoldopam, Fesoterodine, Flibanserin, **Glycopyrronium**, **Homatropine**, **Homatropine methylbromide**, HY10275, Hydroxyzine, Indigotindisulfonic acid, Lofexidine, Loratadine, Loripirazole, Methacholine, **Methantheline**, Methyldopa, Metixene, Mirtazapine, Moxislyte, Naphazoline, OBE101, OPC-28326, Oxymetazoline, Phentolamine, Pilocarpine, Pimavanserin, Pirlindole, Pizotifen, Prazosin, **Propiverine**, Quinidine, Revefenacin, Rociverine, **Scopolamine**, Silodosin, **Solifenacin**, Tegaserod, Terazosin, Terfenadine, Tetryzoline, Thonzylamine, **Tiotropium**, Tizanidine, Tolazoline, **Tolterodine**, Trazodone, **Trihexyphenidyl**, **Umeclidinium**, Xylometazoline |
| 6 | 10 | 75.9% to 83.9% (mean: 78.3%) | Adenosine, Defibrotide, Doxofylline, Dyphylline, Enprofylline, **Etomidate**, Istradefylline, **Midazolam**, Pentoxifylline, Theobromine |
| 7 | 3 | 76.4% to 84.9% (mean: 81.7%) | Clenbuterol, Epinephrine, **XL765** |

**Table 10.** Frequent (≥ 33%) targets of clusters of DrugBank compounds with positive-class likelihood ≥ 75% as estimated by the ensemble trained on male-only datasets.

| Cluster | Target frequency | Target gene name | Target full name |
| --- | --- | --- | --- |
| 1 | 97% | ADRB1 | Beta-1 adrenergic receptor |
|  | 95% | ADRB2 | Beta-2 adrenergic receptor |
|  | 38% | ADRB3 | Beta-3 adrenergic receptor |
| 2 | 84% | DRD2 | D(2) dopamine receptor |
|  | 59% | 5HT2A | 5-hydroxytryptamine receptor 2A |
|  | 47% | 5HT1A | 5-hydroxytryptamine receptor 1A |
|  | 45% | ADA1A | Alpha-1A adrenergic receptor |
|  | 42% | 5HT2C | 5-hydroxytryptamine receptor 2C |
|  | 42% | DRD1 | D(1A) dopamine receptor |
|  | 39% | DRD3 | D(3) dopamine receptor |
|  | 39% | ADA2A | Alpha-2A adrenergic receptor |
|  | 38% | ADA1B | Alpha-1B adrenergic receptor |
|  | 34% | ADA2C | Alpha-2C adrenergic receptor |
|  | 33% | ADA2B | Alpha-2B adrenergic receptor |
| 3 | 42% | KCNH2 | Potassium voltage-gated channel subfamily H member 2 |
|  | 38% | CAC1C | Voltage-dependent L-type calcium channel subunit alpha-1C |
|  | 38% | CALM1 | Calmodulin-1 |
|  | 33% | GBRA1 | Gamma-aminobutyric acid receptor subunit alpha-1 |
|  | 33% | CAC1H | Voltage-dependent T-type calcium channel subunit alpha-1H |
| 4 | 100% | SC6A3 | Sodium-dependent dopamine transporter |
|  | 60% | SC6A4 | Sodium-dependent serotonin transporter |
|  | 60% | VMAT2 | Synaptic vesicular amine transporter |
|  | 60% | ADA2A | Alpha-2A adrenergic receptor |
|  | 40% | HRH1 | Histamine H1 receptor |
|  | 40% | SC6A2 | Sodium-dependent noradrenaline transporter |
|  | 40% | ADA1A | Alpha-1A adrenergic receptor |
|  | 40% | AOFB | Amine oxidase [flavin-containing] B |
|  | 40% | AOFA | Amine oxidase [flavin-containing] A |
| 5 | 41% | ACM3 | Muscarinic acetylcholine receptor M3 |
|  | 41% | ACM1 | Muscarinic acetylcholine receptor M1 |
|  | 41% | ACM2 | Muscarinic acetylcholine receptor M2 |
|  | 39% | ADA1A | Alpha-1A adrenergic receptor |
|  | 36% | ACM4 | Muscarinic acetylcholine receptor M4 |
|  | 36% | ACM5 | Muscarinic acetylcholine receptor M5 |
|  | 35% | ADA2A | Alpha-2A adrenergic receptor |
| 6 | 90% | AA2AR | Adenosine receptor A2a |
|  | 70% | AA1R | Adenosine receptor A1 |
|  | 40% | GBRA1 | Gamma-aminobutyric acid receptor subunit alpha-1 |
| 7 | 67% | ADRB1 | Beta-1 adrenergic receptor |
|  | 67% | ADRB2 | Beta-2 adrenergic receptor |
|  | 67% | TNFA | Tumor necrosis factor |

Cluster 1 consists of drugs targeting Beta-adrenergic receptors (also see Cluster 2 in the mixed-sex classifier results). This is consistent with a top predictive feature “Adrenoceptor family” from the FA InterPro Domains model (Table 3). Most of these drugs are antagonists (drug names ending in -olol, and additionally Bethanidine, Cryptenamine, Labetalol, Propafenone and Sotalol), but some are agonists. As noted above, our models cannot distinguish between activators and inhibitors of the same target, because this information is absent for most compounds in DrugBank and other target databases which we used for training the models. Thus, while these compounds are predicted to have strong effects on murine lifespan, the sign of this effect can be either positive or negative, depending on whether the compound affects the target in the same way as the lifespan-extending molecules in the positive training dataset or in the opposite way. Because we had antagonists of Beta-adrenergic receptors (Metoprolol, Nebivolol ^66^) in our positive training dataset, we predict that antagonists of these receptors will likely extend murine lifespan, whereas the agonists will likely shorten it. Beta-blockers decrease heart rate, thus somewhat supporting the “fixed number of heartbeats per lifespan” hypothesis ^67^. They also decrease cardiac output. In agreement with this, in a recent study of caloric restriction in genetically heterogeneous female mice, cardiac output was negatively correlated with lifespan ^68^. Interestingly, stimulation of ADRB2 by noradrenaline secreted by peripheral sympathetic nerves causes hyperactivation and exhaustion of melanocyte stem cell populations, leading to hair greying ^69^. Encouragingly, a mammalian model of longevity based on interruption of Beta-adrenergic receptor signalling at the level of adenylyl cyclases, specifically disruption of the AC5 gene, has achieved 30% extension of median mouse lifespan^70^. This disruption also protected mice from ageing-induced cardiomyopathy, e.g., hypertrophy, apoptosis, fibrosis, and reduced cardiac function, as well as protected from reduced bone density and susceptibility to fractures^70^. Finally, although invertebrates do not express Beta-adrenergic receptors, they possess a functional analogue of noradrenaline – octopamine^71^. It binds to octr-1, ser-3, ser-6 and several other GPCRs coupled to Gs proteins, which activate adenylyl cyclase and increase cAMP levels, as Beta-adrenergic receptors do in vertebrates. Mianserin, an antagonist of several biogenic amine receptors in nematodes, including octopamine receptors, increased *C. elegans* lifespan by 25-32% ^72,73^. Moreover, recently it has been shown that *C. elegans* lacking the neuronal octr-1 have extended lifespans at warm temperatures but shortened lifespans at cold temperatures^74^. Altogether, this highlights conservation of this pathway across taxa, indicating that it might be as fundamental for regulating longevity as the InsR/IGF-1R pathway.

Cluster 2 consists of drugs targeting primarily Dopamine receptors (also see Cluster 4 in the mixed-sex classifier results); but in addition to that, 5-hydroxytryptamine (Serotonin) receptors and Alpha-adrenergic receptors. However, because we only had a Dopamine receptor agonist precursor (Levodopa ^75^) in our positive training dataset, but no interactors of Serotonin or Alpha-adrenergic receptors, it is likely that Dopamine receptors are driving the effects and clustering of this group of compounds. This is also consistent with a top predictive feature “(Adenylate cyclase-activating) dopamine receptor signalling pathway” from the NE Gene Ontology Process model (Table 4). Drugs from this cluster that act as Dopamine receptor agonists and thus are likely to extend murine lifespan are: Amantadine, Apomorphine, Bromocriptine, Cabergoline, Dihydro-alpha-ergocryptine, Dihydroergotamine, Lisuride, Piribedil, Pramipexole, Quinagolide, Ropinirole, Rotigotine, Sarizotan and Sumanirole. Interestingly, multiple compounds targeting Dopamine receptors, Serotonin receptors and Alpha-adrenergic receptors, including Dihydroergotamine predicted by our model, extended lifespan in *C. elegans* by 10-43% ^73^. This again indicates evolutionary conservation of longevity pathways targeting biogenic amine receptors, which are currently overlooked by the scientific community.

Compounds in Cluster 3 are targeting voltage-dependent potassium and calcium channels, as well as calmodulin and GABA receptor. Several compounds from our positive training dataset (berberine ^76^, chloroquine ^77^, nebivolol ^66^) are listed as interactors of voltage-dependent potassium channels in PHAROS, most likely acting as inhibitors. This is consistent with a top predictive feature “Regulation of membrane potential” from the NE Gene Ontology Process model (Table 4). Additionally, another compound from our positive training dataset, melatonin ^78,79^, is listed as an interactor of calmodulin in DrugBank, and appears to be an inhibitor ^80^. Moreover, Taurine ^81^, another compound from our positive training dataset, is listed in DrugBank as an agonist of GABA receptors. Thus, we predict that compounds inhibiting voltage-dependent potassium channels (Azimilide, Bepridil, Dofetilide) or calmodulin (Bepridil), or activating GABA receptors (Halothane, Isoflurane, Topiramate, Enflurane), are likely to extend murine lifespan. Three inhibitors of calcium channels and two inhibitors of potassium channels extended *C. elegans* lifespan by 15-25% and 12-42%, respectively ^73^. Interestingly, aging has been proposed to be driven by abrogation of the bioelectric signalling that normally harnesses individual cell behaviours toward the creation and upkeep of complex multicellular structures, and that signalling is mediated primarily by ion channels ^82^.

Cluster 4 compounds target Sodium-dependent dopamine, serotonin and noradrenalin transporters, Synaptic vesicular amine transporter, Alpha-adrenergic receptors, Histamine H1 receptor and Amine oxidases. Interestingly, we only had a Histamine H1 receptor antagonist (Meclizine ^83^) and an Amine oxidase B inhibitor (L-deprenyl/Selegiline ^84^) in our positive training dataset, but no interactors of Sodium-dependent dopamine, serotonin and noradrenalin transporters or Alpha-adrenergic receptors. This indicates that compounds in this cluster are not very selective and have multiple targets simultaneously. Nevertheless, we expect drugs inhibiting Histamine H1 receptor (Benzatropine, Diphenylpyraline) or Amine oxidase B (4-Methoxyamphetamine, Metamphetamine) to extend murine lifespan unless the other targets of these drugs will lead to lifespan shortening. Interestingly, it has been shown that harmol, which simultaneously modulates Amine oxidase B and GABA-A receptor, induces mitophagy and AMPK pathway activation, and improves glucose tolerance, liver steatosis and insulin sensitivity in pre-diabetic male mice, delays frailty onset, improves glycemia, exercise performance and strength in two-year-old male and female mice, as well as extends the lifespan of *C. elegans* and female *D. melanogaster* ^85^. Four inhibitors of Sodium-dependent dopamine, serotonin and noradrenalin transporters and three antagonists of Histamine H1 receptor extended *C. elegans* lifespan by 14-34% and 18-32%, respectively ^73^.

Cluster 5 compounds target predominantly Muscarinic acetylcholine receptors and also Alpha-adrenergic receptors. Surprisingly, we did not have any compounds in our positive training dataset that target any of these receptors. Likely, compounds targeting these receptors were prioritised via feature enrichment, especially the NE method. This highlights the ability of our models to go beyond reiterating the targets from the training dataset. Potentially, drugs targeting Muscarinic acetylcholine receptors represent an entirely novel class of lifespan-extending compounds that have not been previously tested in mice. Interestingly, the lack of M3 muscarinic acetylcholine receptors greatly ameliorated impairments in glucose homeostasis and insulin sensitivity in various forms of experimentally or genetically induced obesity in mice ^86^. Moreover, there is a loss of M1 muscarinic acetylcholine receptors in Alzheimer’s disease human brain tissue ^87^. An antagonist of Muscarinic acetylcholine receptors extended *C. elegans* lifespan by 15% ^73^.

Compounds in Cluster 6 target Adenosine receptors and a GABA receptor. As mentioned above, Taurine ^81^ is the only compound from our positive training dataset that interacts with GABA receptors. There are no compounds in our positive training dataset that interact with Adenosine receptors, so similarly to Muscarinic acetylcholine receptors, they were likely prioritised via feature enrichment. This is consistent with a top predictive feature “ADORA2B mediated anti-inflammatory cytokines production” from the NE All categories model (Table 7). Thus, it could be another novel class of lifespan-extending compounds. Adenosine is an immunosuppressive metabolite produced at high levels within the tumour microenvironment. Importantly, adenosine signalling through the A2a receptor expressed on immune cells potently dampens immune responses in inflamed tissues ^88^. Interestingly, compounds targeting Muscarinic acetylcholine receptors, Alpha-adrenergic receptors and Adenosine receptors act as bronchodilators, vasodilators and smooth muscle relaxants. In any case, based on Taurine effects, we predict GABA receptor agonists (Etomidate, Midazolam) to prolong murine lifespan.

Finally, there are only three compounds in Cluster 7: Clenbuterol, Epinephrine and XL765 (Voxtalisib). Clenbuterol and Epinephrine are both agonists of Beta-adrenergic receptors, and because our positive training dataset includes antagonists of Beta-adrenergic receptors (Metoprolol, Nebivolol ^66^), Clenbuterol and Epinephrine are more likely to shorten murine lifespan rather than increase it. XL765 (Voxtalisib) has also been selected in mixed-sex Cluster 1 and is a dual PI3K/mTOR inhibitor which is predicted to increase murine lifespan by analogy with Alpelisib ^89^ and Rapamycin ^90,91^.

Overall, and surprisingly, most predicted longevity drug targets are related to the nervous system function (e.g. various neurotransmitter receptors and transporters and voltage-gated ion channels). Interestingly, it has been shown that extended longevity in humans is associated with a distinct transcriptome signature in the cerebral cortex that is characterized by downregulation of genes related to neural excitation and synaptic function ^92^. Recent studies in model organisms demonstrate that the aging process is frequently modified by an organism’s ability to perceive and respond to changes in its environment. Many well-studied pathways that influence aging involve sensory cells, frequently neurons, that signal to peripheral tissues and promote survival during the presence of stress. Importantly, this activation of stress response pathways is often sufficient to improve health and longevity even in the absence of stress ^93^.

#### 1.7.3 Testing predicted compounds in *C. elegans*

We then decided to test if some of our top candidates can extend the lifespan of the nematode worm *Caenorhabditis elegans.* Although a positive result in nematodes does not guarantee a positive result in mice (e.g. alpha-ketoglutarate, green tea extract, metformin, resveratrol and fisetin worked in the *Caenorhabditis* Intervention Testing Program (CITP^30^) but failed in the (mouse) ITP^29^), and more importantly, a negative result in nematodes does not preclude a positive result in mice (e.g. acarbose, aspirin and meclizine failed in the CITP but succeeded in the ITP)^31^, if the drug works in both species it likely means it affects evolutionally conserved longevity pathways, such as the IGF-1R/IR-PI3K-mTor pathway^32^.

Since we predicted more than 300 compounds, we could not test them all. The first major filter was ensuring that the compound acts on the target in the same way as positive compounds from our training dataset, as extensively discussed in the previous section. For clusters where we did not have a representative compound in the training dataset, we selected the most commonly used drugs targeting the top targets in the cluster. Since worms do not have adrenoceptors but have their functional analogues – octopamine receptors, instead of compounds in male cluster 1 and mixed-sex cluster 2 we decided to test the most commonly used octopamine receptor inhibitors mianserin and epinastine, which were also predicted by our models but for different clusters. We further excluded classified substances, substances not available from major suppliers, overly expensive substances, as well as substances for which the estimated working concentration would be in the range of hundreds or thousands of micromoles. Finally, if the compound has been tested previously in Petrascheck lab and found not to extend lifespan, we did not retest it. Overall, we selected 22 compounds for testing (Table 11). Compounds were applied on day one of adulthood to *C. elegans* in a liquid lifespan assay. 0.5% DMSO was added in control wells.

**Table 11.**
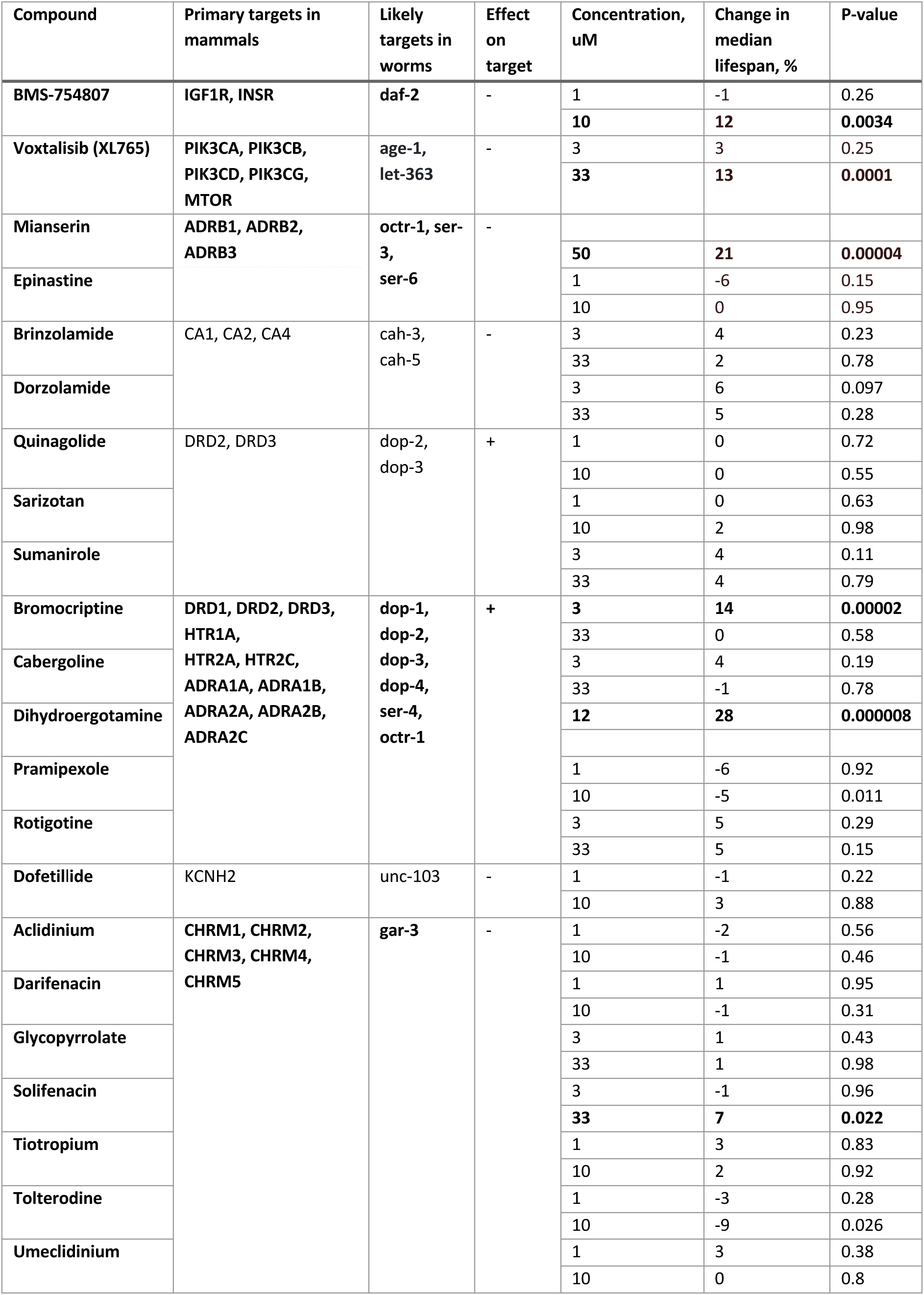
Prioritised compounds tested in *C. elegans* lifespan assay.

Six compounds showed significant extension of median lifespan: dihydroergotamine, mianserin, bromocriptine, voxtalisib, BMS-754807 and solifenacine (Figure 3). Mianserin, a tetracyclic antidepressant used to treat depression and anxiety, has been extensively studied by Petrascheck lab as interacting with octopamine and serotonin receptors and shown to extend *C. elegans* lifespan^72,94,73,95^. Moreover, mianserin is currently undergoing testing in ITP as part of 2024 cohort, with results expected in 2028. Dihydroergotamine, an ergot alkaloid used in the acute treatment of migraine headache and cluster headache, has also been previously shown by Petrascheck lab to extend nematode lifespan^73^. Another ergot alkaloid derivative, bromocriptine, which possesses a potent dopaminergic activity and is used in early Parkinson’s disease, has been shown to extend lifespan in CITP^30,96^. Moreover, lisuride, an ergot derivative that acts as an agonist at dopamine D2 receptors and is used as an anti-Parkinson drug, has been also found to extend worm lifespan as part of a different project that will be published elsewhere. Voxtalisib is a dual PI3K / mTOR inhibitor, currently in trials for the treatment of melanoma, lymphoma, glioblastoma, and breast cancer. BMS-754807 is a potent and reversible IGF-1R / IR inhibitor, currently under investigation in subjects with advanced or metastatic solid tumours. Solifenacine is a commonly prescribed muscarinic antagonist with antispasmodic properties used to treat urinary incontinence. Moreover, we have an indication that umeclidinium, another commonly prescribed muscarinic antagonist used as a long-term maintenance treatment of airflow obstruction in patients with chronic obstructive pulmonary disease, might extend *C. elegans* lifespan at dosages higher than tested here.

**Figure 3.**
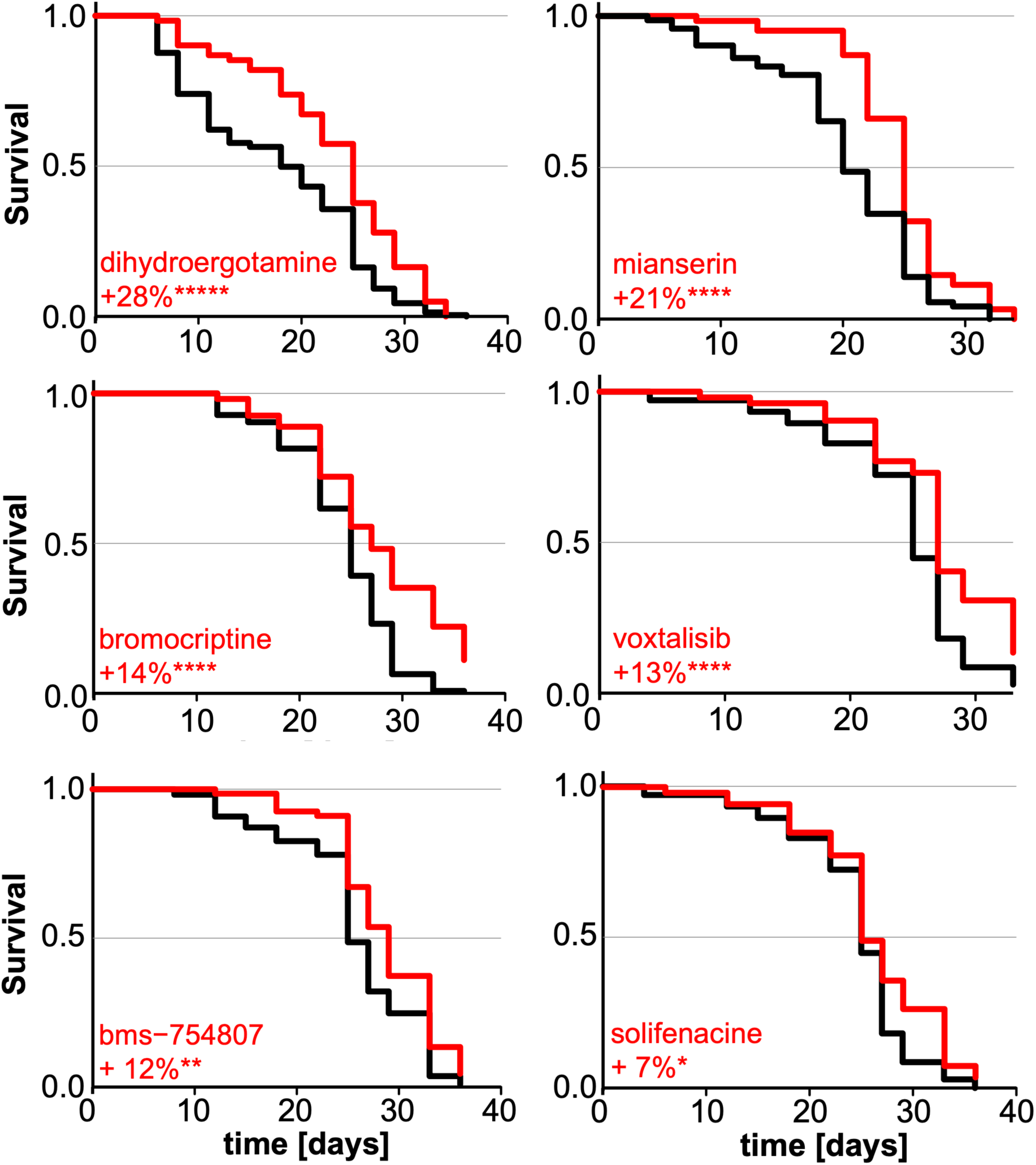
*C. elegans* lifespan assays in liquid medium. Compounds were added on day 1 of adulthood. DMSO 0.5% (black), compound (red). Median lifespan extension in percent is indicated. Survival statistics were calculated using the Mantel–Haenszel log rank test (p-values: * <0.05, ** <0.01, *** < 0.001, **** <0.0001, ***** <0.00001). For technical details see Methods, Table 11 and Supplementary table.xlsx

It can be seen that while some of these compounds target the classical IGF-1R/IR-PI3K-mTor longevity pathway^32^, the others affect targets which were not paid proper attention to by the longevity researchers. Investigating the role of dopamine, serotonin, muscarinic and adrenergic receptors in regulating lifespan might uncover novel highly conserved longevity pathways. One plausible theory is that mianserin and similar molecules acting on dopamine/serotonin receptors interfere with food odour perception, thus inducing behavioural caloric restriction^72,94,56,97^. This fits with our recent finding that pharmacological lifespan extension in male mice strongly correlates with weight loss^23^. Interestingly, male mice have a locus on chromosome 9 which may be modulating longevity through its effect on growth or body weight, but no such locus has been found in females^98^. Top-scoring genes in this locus are Glb1, which is a beta-galactosidase, and Rtp3, which is a receptor-transporting protein that promotes functional cell surface expression of the bitter taste receptors TAS2R16 and TAS2R43. We speculate that higher expression of Rtp3 in some males might make them consume less chow due to more pronounced bitterness and thus reduce weight and promote longevity.

## Conclusions

In this work we report the results of training classification models to predict whether a given chemical compound promotes longevity in mice. Each instance used to train our models was labelled using murine lifespan data from DrugAge, which reflects the current state-of-the-art literature on animal longevity studies. We created datasets using various feature types to describe these compounds, with the most successful models resulting from direct protein target annotations (GO terms, pathways, protein domains and UniProt keywords). Notably, features related to G-protein coupled receptors, especially receptors for neurotransmitters, metabolic hormones and sex hormones, were identified as strong predictors of lifespan extension.

We used the top-performing models to identify compounds with potential for murine lifespan extension, by highlighting consistent false-positive classifications and by creating ensembles to classify over 5000 unlabelled instances from DrugBank (unseen during model training). We clustered the most confident positive-class predictions from the unlabelled data analysis based on their feature similarity, using the DBSCAN clustering algorithm, after applying UMAP dimensionality reduction. Major clusters of prioritised compounds target receptors to IGF1, insulin, adrenaline, noradrenaline, dopamine, serotonin, acetylcholine and adenosine, sodium-dependent dopamine, serotonin and noradrenalin transporters, voltage-gated potassium and calcium channels, as well as carbonic anhydrases. We tested 22 prioritised compounds in the nematode worm *C. elegans* and identified 6 compounds that significantly extend lifespan: dihydroergotamine, mianserin, bromocriptine, voxtalisib, BMS-754807 and solifenacine. This does not preclude other prioritised compounds to be effective in mice.

There are several limitations of our work. First, the number of lifespan experiments performed on *M. musculus* is very small compared to *C. elegans* or *D. melanogaster,* due to their much higher cost and duration, which limits the number of instances available for training the classifiers. This could mean that some important classes of longevity drugs are missing altogether, although our results indicate that our models can nominate novel classes of drugs which were absent in the training dataset, presumably based on shared feature patterns between compounds of different classes. A related limitation is that there are even fewer successful lifespan-extending experiments with female mice, although both sexes are usually tested simultaneously. This could be explained by inherent biological differences in ageing or resistance mechanisms between males and females, as well as potential biases in the choice of compounds for testing. Lack of positive examples limited our ability to train successful classifiers for females; however, we were able to train moderately successful mixed-sex classifiers. Third, for the majority of compounds there is no information in DrugBank and other databases on whether they are inhibiting or activating their targets, which prevented us from constructing datasets and models that can discriminate this property. This led to prediction of multiple compounds with potentially opposing effects on lifespan. However, we were able to at least partially rectify this problem by manually comparing the predicted compounds with those in the training dataset.

In conclusion, this work provides a promising methodology for the preclinical discovery of lifespan-extending compounds in *Mus musculus*, with broader implications for human longevity research. By providing both *in silico* screening tools and biological insights into ageing mechanisms, we pave the way for the development of novel therapeutics targeting ageing and age-related diseases. Future studies should focus on validating the top predictions in mammalian models and exploring the translatability of these findings to humans.

## 2. Acknowledgements

This project was funded by a research grant from the UK’s Biotechnology and Biological Sciences Research Council (BBSRC), grant reference numbers BB/V007971/1 (to AAF and CKF) and BB/V010123/1 (to JPM).

## 3. Author Contributions

JPM and AAF conceived the overall project. CR and AAF designed the machine learning methodology. AVB and JPM designed the target selection and feature enrichment methodologies. AVB and CR created the datasets. CR implemented all machine learning algorithms and ran all the computational experiments. MP performed *C. elegans* lifespan assays. All authors analysed and discussed the results (CR and AAF analysed the results mainly from a machine learning perspective, whilst AVB, CKF, MP and JPM analysed the results mainly from a biomedical perspective). The manuscript was written mainly by AVB and CR, but all authors contributed to writing and revising the manuscript.

## 4 Conflicts of Interest

JPM is CSO of YouthBio Therapeutics, an advisor/consultant for the BOLD Longevity Growth Fund, 199 Biotechnologies, and NOVOS, and the founder of Magellan Science Ltd, a company providing consulting services in longevity science. The other authors declare no conflict of interest.

## 5 Data Availability

The datasets used in the experiments will be made freely available on the web when the paper is published. A public web server with our best classifier ensembles: https://www.cs.kent.ac.uk/projects/lodprime/

## Notes

https://www.cs.kent.ac.uk/projects/lodprime/

### Summary of Updates

Validation in C.elegans by Michael Petraschek was added and he was added as a coauthor, mixed-sex results were transferred to Supplementary, figures and tables added and edited, some other edits to the text

