## Supplementary for "Predicting Mouse Lifespan-Extending Chemical Compounds with Machine Learning"

### Analysis of Feature Importance in the Mixed-Sex Models

Supplementary Tables 1-5 show, for each of our 5 selected models trained on mixed-sex datasets, which 10 features had the highest importance when labelling a compound as positive (predicted to increase lifespan in mice) or negative class.

Notably, for all models trained on mixed-sex datasets, sex is the most important feature, likely because the compounds in our dataset had negative outcomes when tested in female mice twice more often than in male mice, thus the sex variable is highly predictive.

Supplementary Table 1 – Most relevant features in the NE Gene Ontology Components model

| Feature | Description | Class Tendency |
|---|---|---|
| Sex | Sex of the mice used in the experiment | F: Negative class |
| GO:0016323 | Basolateral plasma membrane | Unclear |
| GO:0005834 | Heterotrimeric G-protein complex | Positive class |
| GO:0005886 | Plasma membrane | Positive class |
| GO:1904813 | ficolin-1-rich granule lumen | Negative class |
| GO:0005829 | Cytosol | Negative class |
| GO:0005759 | Mitochondrial matrix | Negative class |
| GO:0000785 | Chromatin | Negative class |
| GO:0120025 | Plasma membrane bounded cell projection | Positive class |
| GO:0005667 | Transcription regulator complex | Negative class |

From Supplementary Table 1 we can see that compounds which target proteins located at the plasma membrane and/or plasma membrane-bounded cell projections, especially the heterotrimeric G-protein complexes, have a high chance of extending murine lifespan. On the other hand, compounds which target proteins located in the cytosol, mitochondrial matrix, chromatin, transcription regulator complex or ficolin-1-rich granule lumen have much lower chances of extending murine lifespan. Ficolin-1 is a crucial protein in the innate immune system, primarily stored in granules within neutrophils. These granules release ficolin-1 into the extracellular environment in response to stimuli, where it binds to carbohydrate structures on pathogens, apoptotic cells, and other particles, thereby activating the lectin pathway of complement activation. The female (F) value of the feature Sex is also associated with the negative class, as in the other tables in this Section.

Supplementary Table 2 – Most relevant features in the NE Wiki Pathways model

| Feature | Description | Class Tendency |
|---|---|---|
| Sex | Sex of the mice used in the experiment | F: Negative class |
| WP536 | Calcium regulation in cardiac cells | Positive class |
| WP3929 | Chemokine signalling pathway | Positive class |
| WP4583 | Biomarkers for urea cycle disorders | Negative class |
| WP399 | Wnt signalling pathway and pluripotency | Negative class |
| WP5046 | NAD metabolism in oncogene-induced senescence and mitochondrial dysfunction-associated senescence | Negative class |
| WP3594 | Circadian rhythm genes | Negative class |
| WP4313 | Ferroptosis | Negative class |
| WP4788 | Autosomal recessive osteopetrosis pathways | Unclear |
| WP5200 | Dravet syndrome | Positive class |

Supplementary Table 2 demonstrates that compounds interacting with proteins involved in calcium regulation in cardiac cells, a chemokine signalling pathway or Dravet syndrome (caused by a loss of function of the voltage-gated sodium channel Nav1.1, affecting the excitability of neurons, particularly inhibitory interneurons) are likely to extend murine lifespan. On the other hand, compounds affecting urea cycle disorders, Wnt signalling pathway and pluripotency, NAD metabolism in oncogene-induced senescence and mitochondrial dysfunction-associated senescence, circadian rhythms, or ferroptosis, are less likely to do so.

Supplementary Table 3 – Most relevant features in the NE Reactome Pathways model

| Feature | Description | Class Tendency |
|---|---|---|
| Sex | Sex of the mice used in the experiment | F: Negative class |
| HSA-9634597 | GPER1 signalling | Positive class |
| HSA-420092 | Glucagon-type ligand receptors | Positive class |
| HSA-1430728 | Metabolism | Unclear |
| HSA-381676 | Glucagon-like Peptide-1 (GLP1) regulates insulin secretion | Positive class |
| HSA-163359 | Glucagon signalling in metabolic regulation | Positive class |
| HSA-5576891 | Cardiac conduction | Positive class |
| HSA-9009391 | Extra-nuclear oestrogen signalling | Positive class |
| HSA-1296071 | Potassium Channels | Positive class |
| HSA-3247509 | Chromatin modifying enzymes | Negative class |

Supplementary Table 3 shows that compounds affecting extra-nuclear oestrogen signalling and G protein-coupled estrogen receptor 1 (GPER1) in particular, glucagon signalling in metabolic regulation, glucagon-type ligand receptors and glucagon-like Peptide-1 (GLP1)-mediated regulation of insulin secretion, as well as cardiac conduction and potassium channels, are predicted to extend lifespan with high probability. Note that both GPER1 and GLP1R are G protein-coupled receptors, making the results of this model in agreement with the results of NE Gene Ontology Components model (Supplementary Table 1) which selected heterotrimeric G-protein complexes as one of its top features. The "Cardiac conduction" feature is consistent with "Calcium regulation in cardiac cells" feature from the NE Wiki Pathways model (Supplementary Table 2). On the other hand, compounds interacting with chromatin-modifying enzymes are less likely to extend murine lifespan according to the predictions. Notably, this is also consistent with the NE Gene Ontology Components model which labelled "chromatin" and "transcription regulator complex" as negative class features (Supplementary Table 1).

Supplementary Table 4 – Most relevant features in the NE KEGG Pathways model

| Feature | Description | Class Tendency |
|---|---|---|
| Sex | Sex of the mice used in the experiment | F: Negative class |
| hsa04929 | GnRH secretion | Positive class |
| hsa05152 | Tuberculosis | Negative class |
| hsa05202 | Transcriptional misregulation in cancer | Negative class |
| hsa04062 | Chemokine signalling pathway | Positive class |
| hsa05017 | Spinocerebellar ataxia | Positive class |
| hsa04022 | cGMP-PKG signalling pathway | Positive class |
| hsa01200 | Carbon metabolism | Negative class |
| hsa05134 | Legionellosis | Negative class |
| hsa05146 | Amoebiasis | Unclear |

The results in Supplementary Table 4 suggest that compounds interacting with proteins involved in gonadotropin-releasing hormone (GnRH) secretion, chemokine signalling pathway, cGMP-PKG signalling pathway or spinocerebellar ataxia are promising candidate lifespan-extending compounds. On the other hand, compounds targeting proteins involved in tuberculosis, transcriptional misregulation in cancer, carbon metabolism or legionellosis are unlikely to be effective for lifespan extension. Please note that chemokine signalling pathway has already been selected as a top feature in the NE Wiki Pathways model (Supplementary Table 2).

Supplementary Table 5 – Most relevant features in the Molecular Fingerprints model

| Feature | Description | Class Tendency |
|---|---|---|
| Sex | Sex of the mice used in the experiment | F: Negative class |
| ≥ 8 H | 8 or more hydrogen atoms | Negative class |
| ≥ 1 N | At least one nitrogen atom | Positive class |
| N(~C)(~C)(~H) | At least one nitrogen atom with two carbon atoms and one hydrogen atom as nearest neighbours, regardless of bond order | Positive class |
| N-H | At least one bonded pair of nitrogen and hydrogen atoms | Positive class |
| C-C-N-C-C | At least one simple SMARTS pattern: C-C-N-C-C | Positive class |
| O=C-C=C-[#1] | At least one simple SMARTS pattern: O=C-C=C-[#1] | Negative class |
| O=C-C-C | At least one simple SMARTS pattern: O=C-C-C | Negative class |
| C=O | At least one pair of carbon and oxygen atoms with a double bond | Negative class |
| C-C-C-O-[#1] | At least one simple SMARTS pattern: C-C-C-O-[#1] | Negative class |

Supplementary Table 5 demonstrates that compounds are more likely to extend lifespan if they have less than 8 hydrogen atoms, at least one nitrogen atom, preferably with two carbon atoms and one hydrogen atom as nearest neighbours, or even better as a C-C-N-C-C pattern. On the other hand, pairs of carbon and oxygen atoms with a double bond, as well as O=C-C=C-[#1], O=C-C-C and C-C-C-O-[#1] patterns, should be avoided.

### Labelling unseen data from an external dataset

Supplementary Figure 1 shows the UMAP projections of the most confident positive-class predictions of the ensemble trained on mixed-sex datasets, as well as the positive class likelihoods of these compounds and their clusters as determined by DBSCAN. Supplementary Table 6 shows the clusters of compounds with positive-class likelihood ≥75% as estimated by the ensemble trained on mixed-sex datasets, and each cluster's most frequent protein targets are shown in Supplementary Table 7.

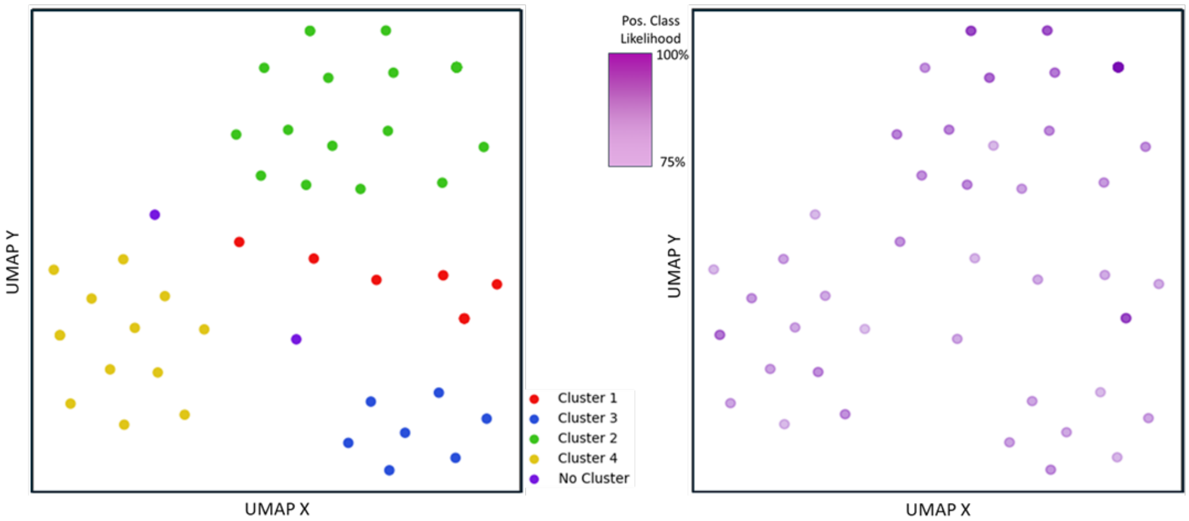


Supplementary Figure 1 – Clustering (left) and positive-class likelihood (right) of DrugBank compounds with positive-class likelihood ≥75% predicted by the ensemble trained on mixed-sex datasets

Supplementary Table 6 – Clusters of DrugBank compounds with positive-class likelihood >75% as estimated by the ensemble trained on mixed-sex datasets. Potential lifespan-extending compounds are highlighted in bold.

| Cluster | Number of compounds | Positive class likelihood Min-Max (Mean) | Compounds |
|---|---|---|---|
| 1 | 8 | 76.4% to 81.5% (mean: 78.3%) | **BMS-754807**, Insulin peglispro, **Linsitinib**, Mecasermin rinfabate, **N-[2-(2-iodo-5-methoxy-1H-indol-3-yl)ethyl]acetamide**, Primaquine, Somatrem, **XL765** |
| 2 | 27 | 75.1% to 90% (mean: 81.8%) | **Acebutolol, Alprenolol, Atenolol, Befunolol, Betaxolol, Bethanidine, Bevantolol, Bisoprolol, Carteolol, Celiprolol, Cryptenamine**, DL-Methylephedrine, Isoetharine, Isoprenaline, **Levobunolol**, Mephentermine, **Metipranolol, Nadolol, Penbutolol,** Phenylpropanolamine, **Pindolol,** Pirbuterol, **Propafenone, Propranolol,** Racepinephrine, **Sotalol, Timolol** |
| 3 | 7 | 76% to 81.1% (mean: 78.5%) | **4-Methylimidazole, Bendroflumethiazide, Brinzolamide, Chlorothiazide, Dorzolamide, Methyclothiazide, n-{2-[4-(aminosulfonyl)phenyl]ethyl}acetamide** |
| 4 | 13 | 75.2% to 82. 1% (mean: 78.6%) | JNJ-37822681, Cinnarizine, **Dihydro-alpha-ergocryptine**, Domperidone, Norclozapine, **Piribedil, Quinagolide**, Rolicyclidine, **Sarizotan**, Sulpiride, **Sumanirole**, Tetrabenazine, Tetrahydropalmatine |
| No Cluster | 2 | 0.762 to 0.782 (mean: 0.772) | Alpha-Benzyl-Aminobenzyl-Phosphonic Acid, LI-301 |

Supplementary Table 7 – Frequent (≥ 33%) targets of clusters of DrugBank compounds with positive-class likelihood ≥ 75% as estimated by the ensemble trained on mixed-sex datasets

| Cluster | Target frequency | Target gene name | Target full name |
|---|---|---|---|
| 1 | 63% | IGF1R | Insulin-like growth factor 1 receptor |
| | 50% | INSR | Insulin receptor |
| 2 | 100% | ADRB1 | Beta-1 adrenergic receptor |
| | 96% | ADRB2 | Beta-2 adrenergic receptor |
| | 37% | ADRB3 | Beta-3 adrenergic receptor |
| 3 | 100% | CA2 | Carbonic anhydrase 2 |
| | 86% | CA1 | Carbonic anhydrase 1 |
| | 57% | CA4 | Carbonic anhydrase 4 |
| 4 | 100% | DRD2 | D(2) dopamine receptor |
| | 54% | DRD3 | D(3) dopamine receptor |

Cluster 1 is a diverse group that includes compounds targeting primarily Insulin-like growth factor 1 receptor and Insulin receptor. Some of them are activators (agonists) of these receptors (Insulin peglispro, Mecasermin rinfabate), while others are inhibitors (antagonists) (BMS-754807, Linsitinib). There are also compounds activating Melatonin receptors (N-[2-(2-iodo-5-methoxy-1H-indol-3-yl)ethyl]acetamide), activating Growth hormone receptor (Somatrem) and inhibiting PI3K/mTOR (XL765). In our positive training dataset, we had inhibitors of IGF-1 receptor (L2-Cmu[1]), PI3K (Alpelisib[2]), mTOR (Rapamycin[3,4]) and an activator of Melatonin receptors (melatonin itself[5,6]). Thus, we can predict that BMS-754807, Linsitinib, XL765 and N-[2-(2-iodo-5-methoxy-1H-indol-3-yl)ethyl]acetamide are likely to increase murine lifespan, whereas Insulin peglispro, Mecasermin rinfabate and Somatrem are likely to shorten it.

Cluster 2 consists of drugs targeting Beta-adrenergic receptors. Most of them are antagonists (drug names ending in -olol, and additionally Bethanidine, Cryptenamine, Propafenone and Sotalol), but some are agonists. Because we had antagonists of Beta-adrenergic receptors (Metoprolol, Nebivolol[7]) in our positive training dataset, we predict that antagonists of these receptors will likely extend murine lifespan, whereas the agonists will likely shorten it.

Cluster 3 compounds are primarily inhibitors of Carbonic anhydrases. In our positive training dataset, we had Butylated hydroxytoluene[8], which is also an inhibitor of carbonic anhydrases. Thus, we predict that most compounds in this cluster are likely to extend murine lifespan. Interestingly, an increase of tissue-specific carbonic anhydrases in mitochondria from middle-aged mouse brain and skeletal muscle has been documented[9]. Moreover, nematodes *C. elegans* exposed to CAH2 have a dose-related shorter lifespan suggesting that high CAH2 levels are life-limiting[9].

Cluster 4 compounds target mostly Dopamine receptors. Most of them inhibit these receptors, but some compounds activate them (Dihydro-alpha-ergocryptine, Piribedil, Quinagolide, Sarizotan and Sumanirole). In our positive training dataset, we had Levodopa[10] which is a Dopamine receptor agonist precursor. Thus, we predict that compounds which activate Dopamine receptors are likely to extend murine lifespan, whereas inhibitors of these receptors will likely shorten it.
